# Modelling the evolution of viral oncogenesis

**DOI:** 10.1101/523373

**Authors:** Carmen Lía Murall, Samuel Alizon

**Affiliations:** Laboratoire MIVEGEC (UMR CNRS 5290, IRD 224, UM), Montpellier, France

## Abstract

Most human oncogenic viruses share several characteristics, such as being DNA viruses, having long (co)evolutionary histories with their hosts and causing either latent or chronic infections. They can reach high prevalences while causing relatively low case mortality, which makes them quite fit according to virulence evolution theory. After analysing the life-histories of DNA oncoviruses, we use a mathematical modelling approach to investigate how the virus life cycle may generate selective pressures favouring or acting against oncogenesis at the within-host or at the between-host level. In particular, we focus on two oncoprotein activities, namely extending cell life expectancy and increasing cell proliferation rate. These have immediate benefits (increasing viral population size) but can be associated with fitness costs at the epidemiological level (increasing recovery rate or risk of cancer) thus creating evolutionary trade-offs. We interpret the results of our nested model in the light of the biological features and identify future perspectives for modelling oncovirus dynamics and evolution.

## Introduction

Understanding viral oncogenicity is traditionally an endeavour of clinical microbiologists and relies on analysing molecular pathways (for a review, see e.g. [23]). Here, we adopt an ecological and evolutionary perspective, which has been extensively applied to study infection virulence over the years [1] and has even re-emerged as a prism through which to analyse cancer dynamics [22].

Few viruses are known to directly cause cancer in humans: Epstein-Barr Virus (EBV), Hepatitis B Virus (HBV), Kaposi’s sarcoma-associated herpesvirus (KSHV), Merkel cell polyomavirus (MCV), Human T-lymphotropic virus (HTLV-1), certain genotypes of Human papillomaviruses (HPVs) and three kinds of polyomaviruses, namely BK virus, JC virus and Simian Virus (SV40). Further details about the oncogenesis and epidemiology of these viruses can be found in other articles in this issue and in that of Chang et al. [5, 13]. Also, the review by Mesri et al. [23] carefully compares the various pathways of viral oncogenes and their roles in triggering the hallmarks of cancer [15].

The evolutionary ecology perspective moves us away from proximate questions of *how* viruses cause cancer towards asking *why* do they cause cancer and, more practically, under what conditions? This approach requires stepping back and looking across human oncoviruses and squinting to look for patterns. Mathematics provides useful tools for this sort of abstraction, especially because stochastic processes [18] or population dynamics feedbacks are difficult to anticipate [19].

In this article, we compare the life cycles of the above mentioned human oncoviruses using variations of classical viral dynamics models [28, 29]. Since a virus that does not transmit from a host is bound to disappear, we develop a ‘nested model’ [24] to consider between-host effects. The model itself relies on a set of ordinary differential equations (ODEs), which we analyse using stochastic simulations that allow for the random evolution of cancer cell populations from infected cells, which we refer to as cancer initiation event [3]. By varying few assumptions and parameters, we can study how oncogenic processes affect virus fitness at the within-host and between-host levels and how these effects depend on the virus life cycle.

### Virulence and viral life cycles

In its most general definition, virulence is the decrease in host fitness due to the infection [31]. In the following, we assume that virus-induced cancer contributes to virulence, even thought its effect on reproductive success can be limited (host mortality may occur late in life and effects on fertility may be unaffected). Figure 1 shows the global prevalence of 6 human oncoviruses as a function of their virulence (the number of yearly worldwide cancer cases divided by their global prevalence). We see that HR-HPVs and HBV are clearly the most clinically important of these viruses as they are responsible for higher numbers of cancer cases. Furthermore, these cancers can happen during host reproductive ages. Meanwhile, EBV and MCV are the human oncoviruses with the lowest virulence as they are nearly ubiquitous yet are responsible for less than half the number of cancer cases per year. We exclude from our analysis other polyomaviruses and non-HR HPVs given poor global estimates of viral prevalence. Note that the upper right corner of this graph is empty. In animal viruses, we know of at least one example, Marek’s Disease Virus (MDV), a virus from the *Herpesviridae* family that was both prevalent and fairly virulent before vaccination [32]. This shows that large DNA oncoviruses can reach high levels of virulence. The observation that no human oncovirus with high carcinogenicity reaches high prevalence reinforces our assumption that cancer contributes to virulence and is selected against in viral population.

**Figure 1:**
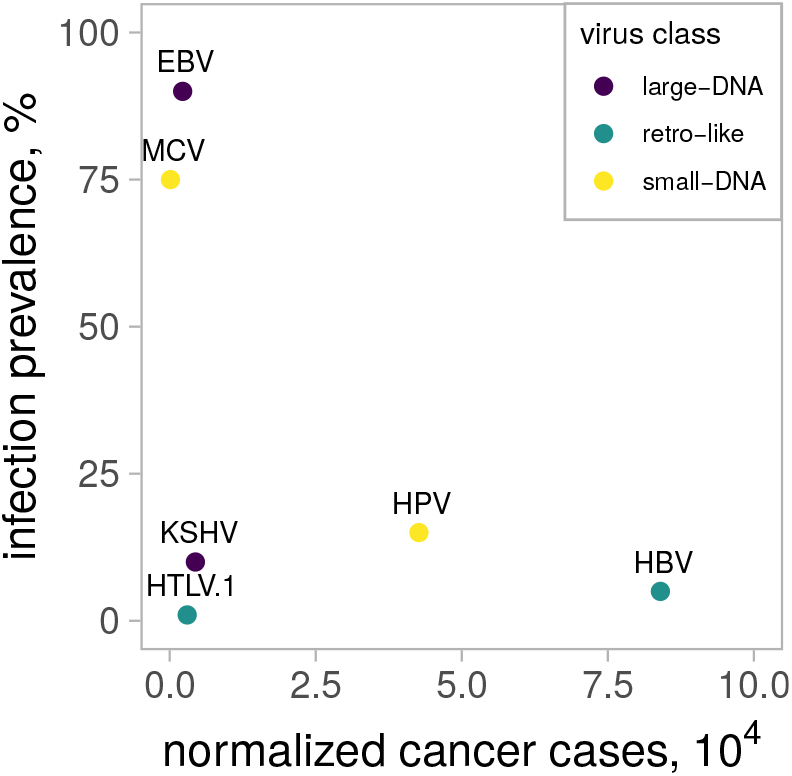
Global prevalence and virulence of oncovirus infections in humans. High-risk HPV and HBV are less prevalent globally but cause more cancer cases per infection. In contrast EBV and MCV are very prevalent but cause fewer cancer cases per infection. The x-axis is obtained by dividing the estimated number of cancer cases worldwide by the global prevalence. Data originates from [21].

Human oncoviruses are not monophyletic which implies that oncogenic traits evolved separately in a case of convergent evolution. Even in the case of HPVs, the two types responsible for the majority of cancers globally, HPV16 and HPV18, belong to different HPV species and the other genotypes in their species are significantly less oncogenic [4, 35]. Oncoviruses can be classified based on their genetic make up and replication modes. For instance, HPVs and polyomaviruses belong to the small DNA viruses, whereas EBV and KSHV belong to the large DNA viruses. In Table 1 we summarise key features and traits of these human oncoviruses, in order to illustrate how we abstracted their life cycles into three distinct groups. As seen in the table under ‘viral class’, we chose to group HBV and HTLV-1 as ‘retro-like’ viruses because HBV is a DNA virus with retrovirus features, while HTLV-1 is a true RNA retrovirus.

**Table 1:** Human oncovirus properties. Adaptation and extension of Table 1 from [21].

|  | EBV | KSHV | HR-HPV | nonHR-HPV | MCV | JCV, BKV, SV40 | HBV | HTLV-1 |
| --- | --- | --- | --- | --- | --- | --- | --- | --- |
| *Cancer class* | 1 | 1 | 1 | 2A,2B,3 | 2A | 2B | 1 | 1 |
| *Viral class* | large dsDNA | large dsDNA | small dsDNA | small dsDNA | small dsDNA | small dsDNA | ss and dsDNA (with retrovirus features) | (+)ssRNA retrovirus |
| *Cell tropism* | lymphocytes, epithelium | endothelium, B-cells | epithelium | epithelium | Merkel cells, epithelium, kidney/bladder | several cell types, epithelium, kidney, CNS | hepatocytes | lymphocytes |
| *Acute/Chronic* | chronic/persistent | chronic/persistent | mainly acute (some persist) | mainly acute | chronic | chronic | acute (chronic if in childhood) | chronic |
| *Lytic/Non-lytic* | latent <> lytic | latent <> lytic (mostly in latent phase) | non-lytic, non-budding | non-lytic, non-budding | lytic (no latency found yet) | lytic and latent | non-lytic, budding | non-lytic, budding |
| *Transmission route* | saliva, perinatal | saliva, blood, sex contact | sex contact, skin contact | sex contact, skin contact | skin contact, perinatal | faecal-oral, sex contact | perinatal, sex contact, blood | perinatal, blood |
| *Oncogenes* | EBNA, LMP1, LMP2A, miRNAs | LANA 1, KSHV-encoded cyclin | E6, E7, E5 |  | LT | LT and sT | HBx | Tax |
| *Oncogene(s) activities* | cell proliferation, cell survival, control cell differentiation | cell proliferation, cell survival, immune evasion, angiogenesis, tether episomes | cell proliferation, cell survival, immune evasion, cell-cycle re-entry, prevent apoptosis |  | pRb binding | cell proliferation, prevent apoptosis (for BK: interfering with p53 & pRb) | cell proliferation, cell survival, immune evasion, cell-cycle re-entry, prevent apoptosis | cell proliferation, cell survival, re-entry |

Summarising the properties of human oncoviruses highlights some common features (although there are exceptions). On average, human oncoviruses are mostly DNA viruses (HTLV being the exception) with a tropism for epithelium-related cells and immune cells. Most of them cause, or at least can cause, chronic infections. Their genomes contain oncogenes that increase the proliferation and survival rate of their host cells (though they can have many more functions). The highest degree of variation between these viruses comes from their transmission routes (even though they all involve close contact and bodily fluids) and their life cycles (with or without a latent stage and lytic or non-lytic). This is why our model focuses on the importance of the latter.

All of these viruses can only persist in human populations over the long-term through between-host transmission. Therefore, unless there is vertical transmission, the fittest virus strains are the ones that maximise infection duration, while maintaining the production of enough infectious viral particles. This clearly leads to trade-offs. For instance, mechanisms such as immune escape or immunosuppression that can decrease host recovery rate are also associated with cancers [11]. Similarly, increased production of virus particles can simultaneously increase transmission rate and virulence, as observed in the case of HIV [12]. Finally, it has also been argued that increased viral replication could lead to more rapid host recovery, e.g. in the case of HPV [26]. Overall, the fittest virus at the within-host level (i.e. the one that infects the highest number of cells) is not necessary the one causing the highest number of secondary infections. To investigate this conflict between levels of adaptation, we resort to mathematical modelling.

Capturing the dynamics of different oncoviruses requires different models. However, in order to identify the effect of life cycle properties such as latency or budding on the fitness of an infection, we need a basis for comparison. We, therefore, only vary the structure of the life cycle itself and homogenise parameter values for our three classes of viruses. This has a cost in terms of biological realism (e.g. our ‘large DNA’ virus is idealised and will not correspond exactly to EBV or KHSV) but it allows us to better understand the selective constraints acting on oncogene activity and to see whether life cycle properties are sufficient to recover known differences in infection phenotypes such as duration or cancer risk.

## Models

To abstract the various life cycles of these oncoviruses, we start with a model that is generic enough to represent any target cell population. Figure 2A shows an uninfected cell population with cells at rest (in phase G_0_ of their cycle), denoted *G_u_*, and those that are in the replication phases of the cell cycle (namely G_1_, S, G_2_ and M), denoted *R_u_.* Each cell division event results in two daughter cells at rest at a rate *2**δ***. The cells at rest die naturally at a rate *μ* and enter replication at a rate **σ**.

**Figure 2:**
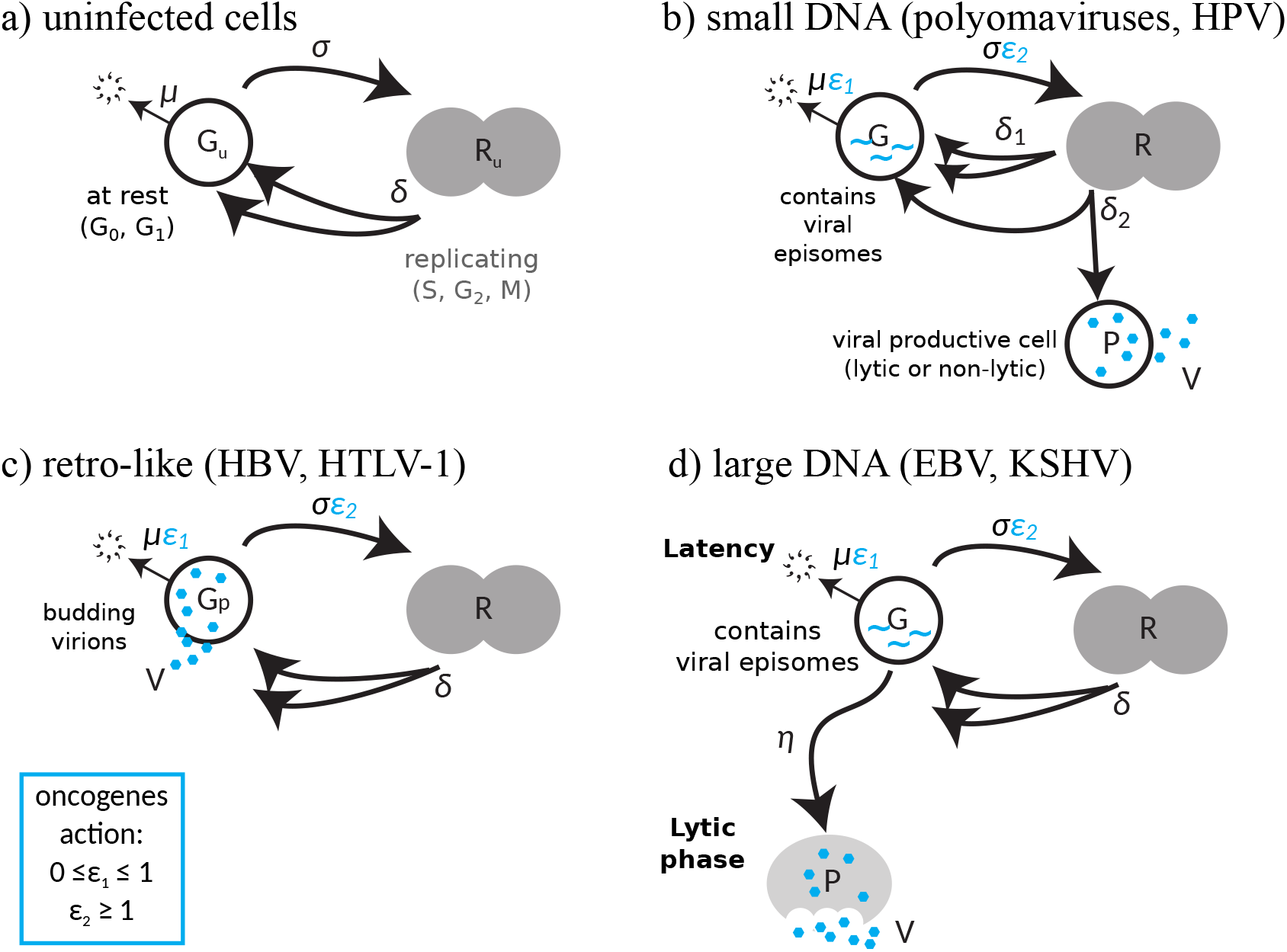
Human oncovirus life cycles. (a) For uninfected cells, generic host cells (*G_u_*) enter the replication phases of the cell cycle (*R_u_*) and produce two daughter cells. (b) In small DNA viruses, infected cells that contain virus in episomal form (*G*) divide and produce either two similar daughter cells or makes virus-producing cells (*P*) which can be lytic, i.e. kills the cells during viral production (e.g. polyomaviruses, such as MCV) or non-lytic, i.e. cells die at natural death rate (e.g. HPVs) and then releases virions (*V*). (c) For retro-like viruses, infected cells (*G_p_*) produce new virus particles (*V*) that bud out from the cell’s membrane. (d) For large DNA viruses, infected cells mostly exist in latent phases with the virus in episomal form (*G*) and virus producing infected cells (*P*) are only made sporadically when a lytic phase is activated, which happens at rate *η*. Oncogenes lengthen cell life (*ε*_1_) and increase cell divisions (*ε*_2_).

Our infection models capture two main activities that viral oncoproteins share across human oncoviruses and that affect the hallmarks of cancer (Table 1 and [23]): extending cell life expectancy (e.g. resisting cell death by preventing cell apoptosis) and increasing cell proliferation and sustaining a proliferative program (e.g. inactivating the G_1_/S checkpoint and other tumour suppressor checkpoints and targeting RB1 and p53). These correspond to parameters *ε*_1_ and *ε*_2_ respectively.

In the models with infections, new cell classes appear such as cells infected with viral episomes (*G*), virion-productive cells (*P*), or budding cells (*G_P_*). The density of free virions is denoted *V*. The flow diagrams in Figure 2 illustrate the mathematical models for each oncovirus group and the equations can be found in the Supplementary Information along with the R scripts used for the simulations. Practically, we implemented stochastic simulations of this system of ODEs using the *τ*-leap Gillespie algorithm [14]. The assumptions related to this implementation are further described in the Supplementary Information and in the Discussion. The simulations allow us to store the number of divisions a cell has been through. Therefore, *G* is the sum of populations of cells *G_D_*, where *D ∈ ℕ* is the number of divisions the cell has gone through.

Although not shown in Figure 2, we introduce a population of cytotoxic T-cells (CTLs), T, in each of the models following many previous models [2]. Without these, virus populations would grow exponentially. Instead, we observe a wider range of immunological scenarios. To avoid unrealistic population densities, we assume a carrying capacity for the total number of infected cells and the total number of CTLs.

We also include a stochastic ‘catastrophic’ event, which corresponds to a cancer initiation event. The rate at which this event occurs at time *t* is given by *v* Σ*_D_ G_D_D^p^*, where *v* is a normalising constant parameter, D is the number of divisions a cell from the population *G_D_* has been through, and *p* is parameter capturing the increase in cancer risk with the number of cell divisions. The rationale behind this assumption is that the lifetime risk of cancer in a tissue correlates with the number of stem cell divisions in the lifetime of this tissue [34]. By default, we assume that *p* = 1 but explore non-linear relationships in Supplementary Results.

Finally, to model the between-host level, we assume that infected hosts regularly interact with other hosts. Upon contact, the virus is transmitted with a probability that depends on the number of free virions (*V*) at that time. Based on epidemiological data from HPV [36], we assume the probability of transmission saturates rapidly with increasing virus load.

Within-host simulations are run until one of the three possible outcomes of the model is reached: the virus is cleared, a cancer initiation event occurs, or the maximum time (set to 50 years) is reached. All the parameter estimates and initial conditions that are similar across all three viral groups were kept constant to facilitate the comparisons between life cycles. These are shown in Supplementary Information and chosen to be biologically realistic (see [27] for more details).

## Results

### Infection kinetics

Figure 3 illustrates typical time series for each of the three infection models. Each plot shows three stochastic realisations of the same model with the same parameter values. The three panels show three combinations of oncogene activity. In Figure 3a, the oncogenes have no action on the death rate and the replication rate of the infected cell (*ε*_1_ = 0 and *ε*_2_ = 1). For two of the life cycles (large-DNA and retro-like), we observe persistent infections, whereas for the small-DNA life cycle the infection is cleared. This make sense given that in the first two life cycles, the homoeostasis parameters of the uninfected cells are barely affected by the infection (there is some increased cell death due to the immune response but it is compensated by rare reinfections by free virions). Notice also that that the stochasticity is strong in these dynamics.

**Figure 3:**
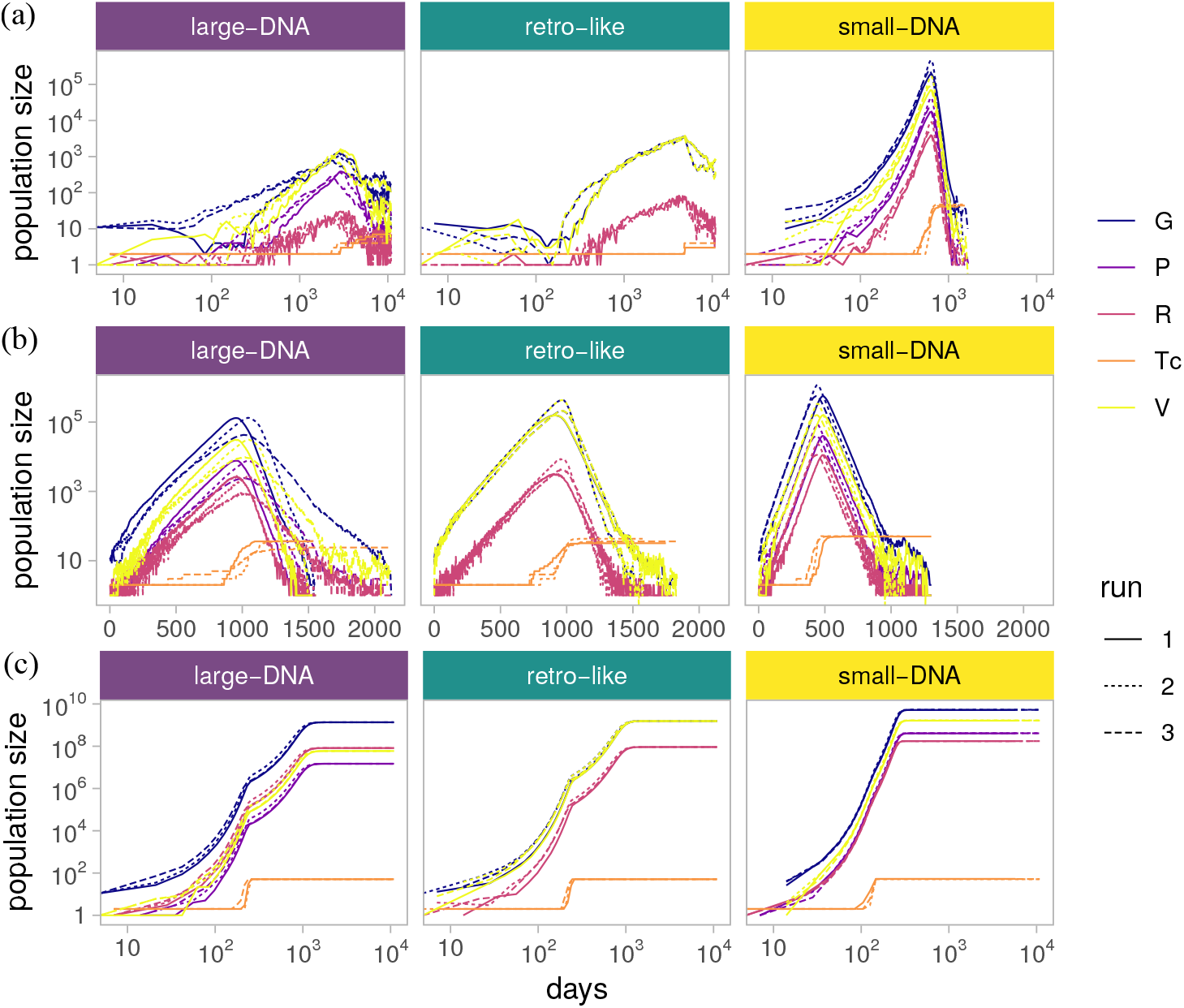
Example within-host population dynamics with (a) no, (b) limited and (c) strong action of the oncogenes. In a), *ε*_1_ = 0 and *ε*_2_ = 1, in b) *ε*_1_ = 0.5 and *ε*_2_ = 1 and in c) *ε*_1_ = 0.5 and *ε*_2_ = 3.5. Each line type corresponds to a stochastic run. Colours indicate infected cells at rest (*G*), infected cells dividing (*R*), infected cells producing virions (*P*), virions (*V*) and cytotoxic T-cells (*T*). For clarity, time is shown in a log scale for persisting infections (in c). Other parameter values are default (see Supplementary Information).

In Figure 3b, we set *ε*_1_ = 0.5, while keeping the other parameters unchanged. This increase in oncogene activity perturbs homeostasis, with a more rapid increase in the immune response (in orange), such that all life cycles lead to clearance. Finally, in Figure 3c, oncogenes increase the infected cell’s life-expectancy and replication rate (*ε*_1_ = 0.5 and *ε*_2_ = 3.5). This allows populations of infected cells to avoid clearance and achieve long-term persistence at high densities. As we will see below, cancer initiation events are only observed in such persistent infections.

### Oncovirus fitness landscapes

We then explored 984 different combinations of (*ε*_1_, *ε*_2_) and performed 25 stochastic simulations per combination. Figure 4a shows the mean duration of the infection. For large-DNA and retro-like viruses, starting from the bottom left corner, increasing oncogene activity, either by decreasing the death rate of the infected cell (*ε*_1_) or increasing its proliferation rate (*ε*_2_), decreases the duration of the infection. Increasing further oncogene effect increases infection duration, with *ε*_2_ having the largest effect. Finally infection duration decreases again for very large values of *ε*_1_ and *ε*_2_. For small-DNA viruses, increasing oncogene action increases infection duration right away. As for the other life cycles, higher values of *ε*_2_ decrease infection duration.

**Figure 4:**
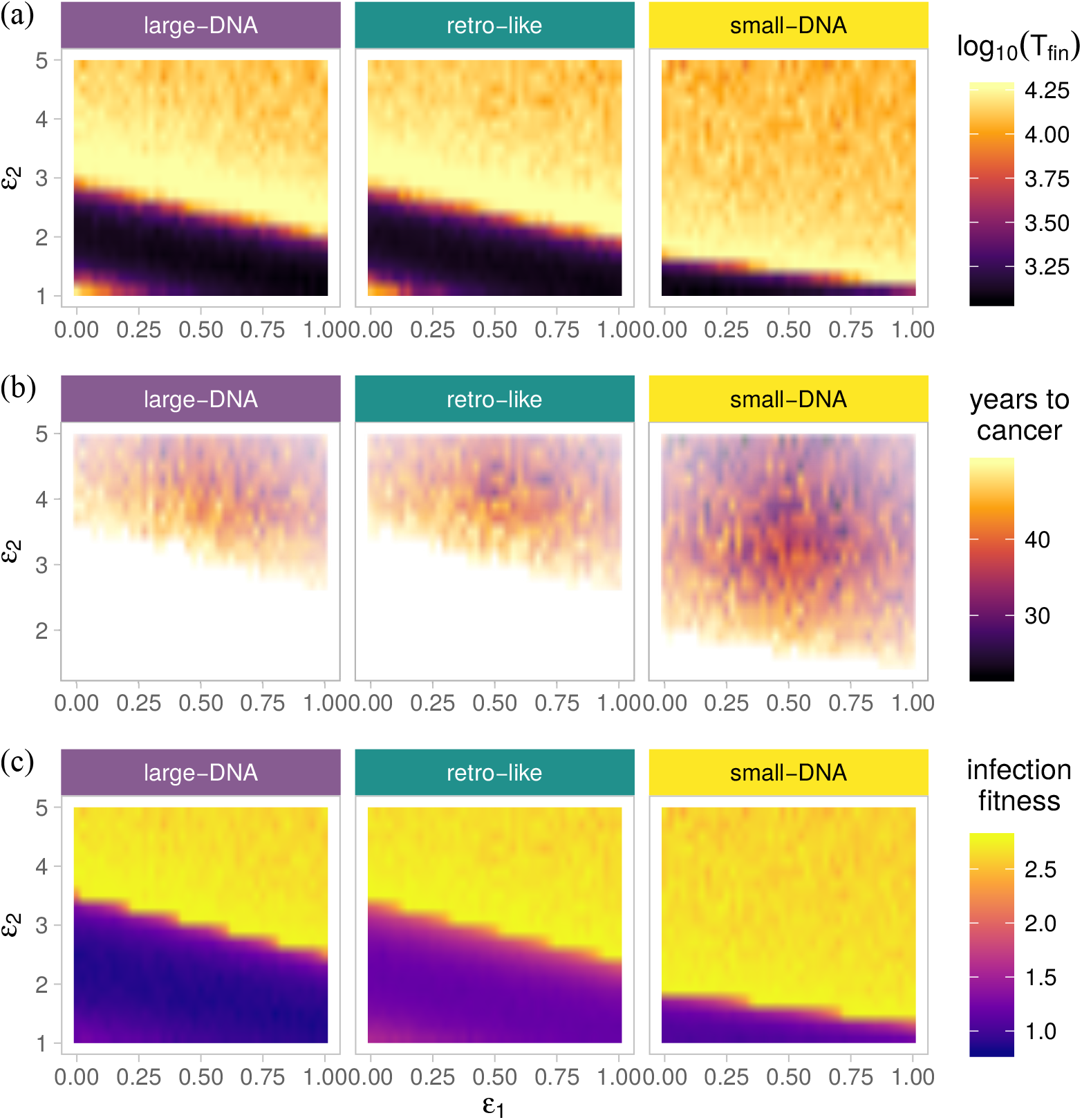
Oncogene effects in the three virus classes on (a) infection duration, i.e. mean final time reached, (b) time to cancer and (c) infection fitness. For each parameter set, the time to cancer is calculated as the average of the time until a cancer initiation event for all the simulations, where such an event occurred. In white, cancer is never observed. Infection fitness is the log_10_ of the number of potential transmission events. We explored 984 combinations of (*ε*_1_, *ε*_2_) with 25 simulations per combination. Other parameter values are default (see Supplementary Information).

Figure 4b shows the average time until a cancer initiation event for parameter sets where such an event occurred. For a given parameter set, the fraction of the 25 simulations that lead to cancer is negatively correlated with the time until this event occurs (Figure not shown). That cancer occurs more rapidly for stronger activity of the oncogenes coincides with the decrease in infection duration reported above. We again see a difference between the large-DNA and retro-like life cycles, for which most of the parameter sets studied do not lead to cancer (the white areas), and the small-DNA viruses, where cancer occurs more often. *ε*_2_ has the strongest effect on the occurrence of cancer initiation events and their timing, but *ε*_1_ also matters, especially for large-DNA and retro-like life cycles, as indicated by the slope separating the white area from the coloured one. Note that here we assumed that the number of divisions an infected cell has been through increases cancer risk linearly. If this risk is independent of the number of divisions or, conversely, if it increases more than linearly, we observed a similar shape but the cancer initiation event occurs more rapidly (Supplementary Figure S2).

Finally, in Figure 4c we show the infection fitness, that is the number of potential secondary infections (see the Supplementary Information). This measure combines infection duration and virion production, which monotonically increases with oncogene activity (Figure not shown). The pattern strongly resembles that in Figure 4a, suggesting that infection duration is the most important component of the infection for virus transmission. Oncovirus fitness is maximised for intermediate values of *ε*_2_. The optimal value of *ε*_1_ depends on the value of *ε*_2_, although to a lesser extent for the small-DNA life cycle. Large-DNA and retro-like viruses exhibit a local fitness peak in the area with very limited action of the oncogenes but its height is limited because few virions are produced (see Figure 3a).

### Maximising fitness

Most of the 984 parameter sets we explored lead to infections that produce virions and last long enough to be transmitted to other hosts. However, variations in infection fitness are such that viruses bearing non-optimal traits are likely to be rarely detected. In Figure 5, we analyse the properties of the ‘fittest’ viruses, that is the parameter sets that lead to the highest between-host fitness values in Figure 4c. In practice, for each life cycle we selected the 25 parameters sets with the highest fitness values and without any cancer initiation event. We then did the same for parameter sets with a cancer initiation event. Our goal was to compare oncovirus strategies that avoid cancer to those that do not.

**Figure 5:**
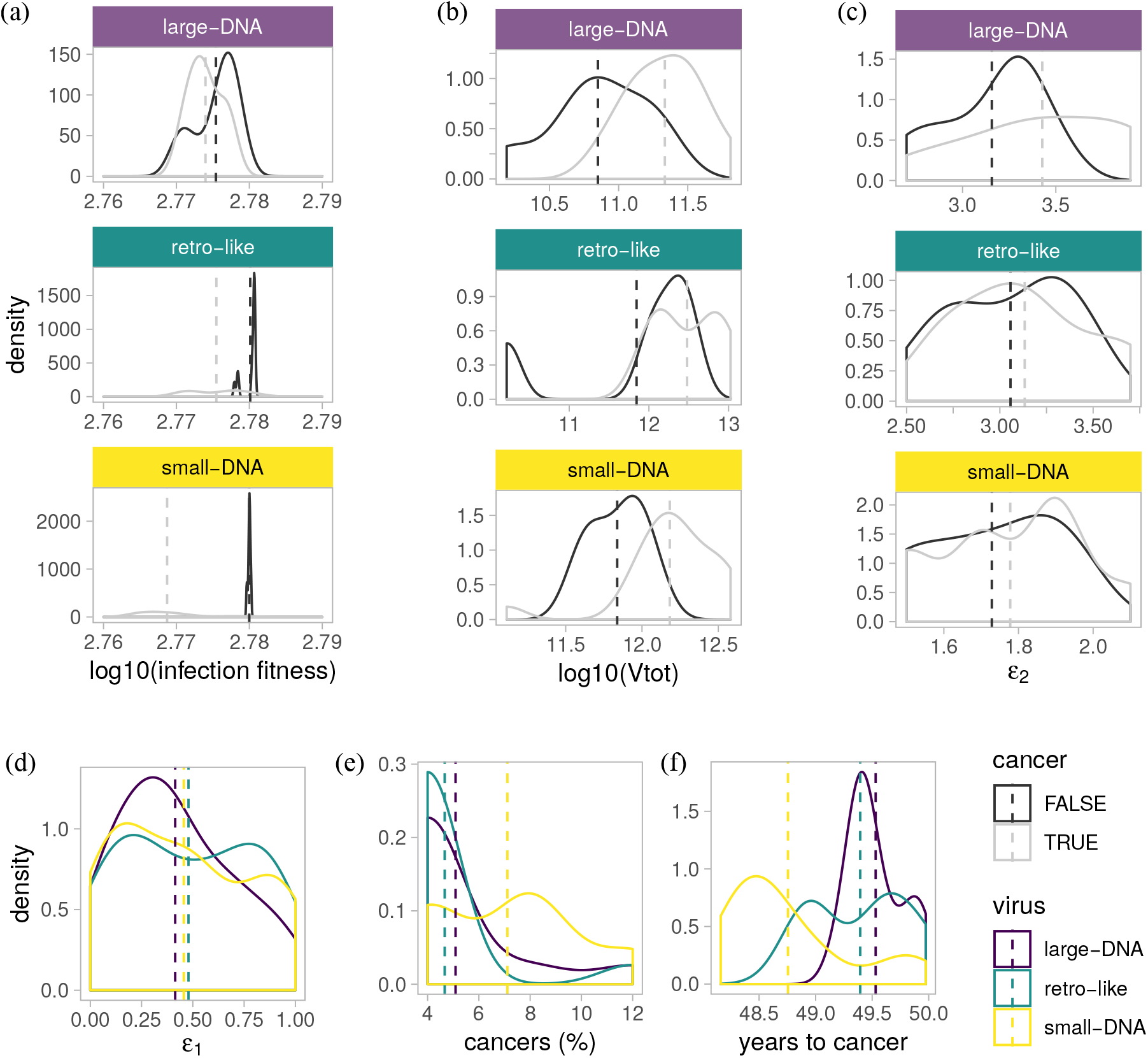
Characteristics of infections maximising virus between-host fitness. a) Infection fitness, b) total number of virions produced during the infection, c) *ε*_2_, d) *ε*_1_, e) for each parameter set, the fraction of the simulations that lead to cancer and f) the years until cancer initiation event. In a, b and c, the 25 fittest sets with cancer are in grey and the 25 without cancer are in black. Dashed line show mean values.

In Figure 5 an, we see that for the fittest parameter sets, infection fitness is higher when there is no cancer (in black) compared to when cancer occurs (in grey). However, these differences are small, especially for the large-DNA life cycle. Interestingly, at the within-host level, the sets with cancer occurrence yield higher total virion production over the course of the infection (Figure 5b). This illustrates the importance of selection as a multi-level process, where a strategy maximising fitness at the within-host level may not be the fittest at the between-host level.

Next, we focus on the oncogene activity associated with these fittest parameter sets. As expected, in Figure 4c we find that the increase in replication rate, *ε*_2_, is much lower for small-DNA viruses. We also find that sets with cancer exhibit on average higher values of *ε*_2_. For *ε*_1_, we find no consistent different in runs with or without cancer (Figure not shown) but we find its value to be slightly larger in retro-like life cycle (Figure 5d). This could be due to its life cycle, where the *G* stage is the one producing the virions, thus, increasing the life-expectancy of the cell also increases the time is spends in the virion-producing stage.

Intuitively, we might have expected a strong effect of *ε*_1_ in the large-DNA virus life cycle because it can take years before a *G* cell becomes a virion-producing cell (*P*). This absence of effect is a consequence of the assumption of the τ-leap Gillespie algorithm, which is memory-less (the time spent by a single cell in a *G* state does not affect its probability to switch to a *P* state). In the Supplementary Information, we use a classical modelling technique [20, 6] to prevent *P* cells from being produced too quickly ‘by chance’ (Supplementary Figure S3). This leads to a decrease in infection fitness, a strong increase in *ε*_1_, and a limited increase in *ε*_2_ in the fittest parameter sets for the large-DNA life cycle.

Finally, for the fittest parameter sets with cancer, we show the fraction of simulations where cancer events occurred (Figure 4e) and the average time at which it occurred (Figure 4f). The small-DNA life cycle stands out with more cancers occurring more rapidly. That large DNA viruses tend to cause cancer later than the other two groups is consistent with the biology (Table 1).

## Discussion

Cancer is often presented as an evolutionary dead-end. For the host, it clearly bears little adaptive value. For oncoviruses, the problem is less straightforward. One obvious cost is that the host may die. Another direct cost is that infected cancer cells frequently contain incomplete viral genomes and tend to produce less or no virions. However, oncogenes have pleiotropic effects (often long before cancer appears) and can be associated with increased virus fitness (e.g. if they increase cell replication rate, thereby increasing virus load, or cell life expectancy).

In the end, the optimal level of viral oncogenesis is likely to result from the balance between selection at the within-host level for increased virus load and at the between-host level for infection duration. This conflict is particularly clear when analysing the fittest virus strategies: the strategies that can lead to cancer have higher virus production than the ones where cancer never occurs, but the latter are associated with more between-host transmission. This is because infection duration largely governs epidemiological fitness, which, in our study, likely results from our assumption that the probability of virus transmission per contact is high (in the case of HPV it has been estimated to be close to 90% [36]). With high infectivity per contact, virus load matters less than the number and frequency of these contacts when it comes to between-host fitness. However, infection duration may not be independent from virus load. For instance, HR-HPV chronic infections that regress tend to exhibit decreasing virus loads, whereas those that persist have constant or increasing virus loads [9].

In our modelling approach, we have voluntarily restricted the range of scenarios to explore by assuming that viruses from the three life cycles have the same parameter values. In spite of this, we find differences between the life cycles that are consistent with the biology. Perhaps the most striking difference has to do with the fitness landscape. Indeed, for the large-DNA and retro-like life cycles, the virus can achieve long-term persistence even with very limited action of the oncogenes. For the small-DNA life cycle this is not possible and we always have strong selection for increased replication rate of infected cells. We also find cancers to be more frequent in small-DNA viruses but they appear later in large-DNA viruses. Another interesting insight of our model comes from the differential effect of extending cell life duration, in that it depends on the virus life cycle. Indeed, for retro-like viruses spending more time in the *G* stage is interesting because this is the virion-producing stage. For large-DNA viruses, the strong added value of decreasing the death rate of the infected cell is only apparent if we add memory into the Gillespie algorithm (Supplementary Figures S3 and S4).

Many evolutionary biology models nest within-host dynamics into an epidemiological framework [24]. The most delicate step in these models is the linking between within-host variables (e.g. virus load, number of target cells, number of immune cells) and epidemiological parameters (such as virulence, transmission rate and recovery rate). Transmission rate can be safely assumed to be related to virus load, but predictions are more difficult when it comes to virulence. Indeed, in experiments this trait is measured using *ad hoc* proxies (e.g. anaemia, decrease in body mass, case fatality ratio, time to death) even though theory predicts qualitative differences in virulence evolution depending on the measure used [8]. If virulence corresponds to cancer and if this event is explicitly excluded into the within-host model, the nesting becomes more intuitive and has a stronger mechanistic basis than with other viruses.

There are several ways in which our model could be extended. For simplicity, we stopped our simulations after the cancer initiation event event. However, our model allows us to follow the fate of a single cell that has become carcinogenic and there is now a wealth of mathematical models to lean on [10, 3]. A possibility for future work would be to include stochasticity in the within-host spread of cancer clones. As discussed elsewhere [16], the within-host environment at the time of cancer emergence, especially the activation state of the immune response, is likely to govern the probability of fixation. Further, we have not included interference of oncoproteins with the immune response through immunosuppression or immune evasion. These would be particularly interesting because they would affect the age distribution of the population of infected cells (see Supplementary Figure S1). Since the number of divisions a cell has been through increases cancer risk [34], this oncogene action would add a mechanistic link to cancer occurrence). In general, our modelling of the immune response is obviously a great simplification of the reality. However, similar assumptions are commonly used when modelling virus dynamics [37, 7, 33] and here it was especially important to be able to compare the three virus life cycles. Finally, another aspect we simplified has to do with the structure of the host tissue. Indeed, for viruses infecting tissues with 3D structures, such as epithelia, this structure could directly impact infection duration [27]. However, this would require virus-specific models since oncoviruses infect different tissues.

Here, we aimed to begin a theoretically grounded conversation about the evolution of viral oncogenesis. We, thus, end with various lines of inquiry. For instance, we considered that variations in oncogenicity can, in part, be explained by virus genetics. For DNA viruses, this could be challenged given the large variation observed between patients and the low viral evolutionary rates. However, recent evidence shows that even when considering only one genotype, HPV16, the E7 gene exhibit less variability in samples from pre-cancers/cancers compared to the controls [25]. Similarly, one could also question whether the viruses we see are the fittest. Indeed, there could be physical constraints preventing the virus from reaching parts of the parameter space. In addition, the coevolutionary dynamics between humans and DNA oncoviruses could also be non-equilibrium processes. In the case of HPV16, it has recently been argued that the most virulent lineage known currently (HPV16A) could originate from Neanderthals and therefore be less adapted to modern populations [30]. Overall, understanding the evolutionary constrains and conditions that explain varying degrees of viral oncogenicity should be studied more widely. Not only for academic interest but also for gleaning new insights into how to design evolution-proof [17] intervention strategies.

## Supporting information

Supplementary methods and results

summary statistics extracted from the simulation data

R scripts to resimulate the data and generate the plots

## Acknowledgments

This work was supported by the European Research Council (ERC) under the European Union’s Horizon 2020 research and innovation program (EVOLPROOF, grant agreement No 648963). The authors acknowledge further support from the CNRS, the IRD and the itrop HPC (South Green Platform) at IRD montpellier, which provided HPC resources that contributed to the results reported here (https://bioinfo.ird.fr/).

## Supplementary Information

One PDF file containing Supplementary Methods and Supplementary Results and two files with R scripts and simulation data.

