## Supplementary methods and results for "Modelling the evolution of viral oncogenesis"

### Supplementary Information to
### *Modelling the evolution of viral oncogenesis*

Carmen Lía Murall and Samuel Alizon

### S1 Supplementary Methods

#### S1.1 Within-host dynamics models

##### S1.1.1 Uninfected cell model

The underlying model used to represent a population of target cells follows the same form for all our viral infections models. A generic population of target cells is broken up into two classes, one of resting cells, $G$ (in phase $G_0$), and cells that are dividing (in phases $G_1$, S, $G_2$, or M), $R$. Cellular dynamics are captured by the following set of ordinary differential equations (ODEs):

$$\dot{G_U} = 2\,\delta\,R_U - (\sigma + \mu)\,G_U \tag{S1a}$$

$$\dot{R_U} = \sigma\,G_U - \delta\,R_U \tag{S1b}$$

Uninfected resting cells, $G_U$, enter the cell cycle and move to a state, $R_U$, at rate $\sigma$. We assume cells enter cell division every 50 days (this of course varies between human cell types, e.g. some epithelial cells have a life cycle of 33 days [3] and memory or naive T cells can live for longer than a year) [9, 1]. Once the cycle is completed, the two daughter cells enter the resting cell class at a rate of $\delta$. Note that the average human cell takes approximately 24 hours to complete cell division and return to resting [7]. Finally, cells at rest have a natural death rate of $\mu$. For the cell population to be at homeostasis, i.e. remain constant over time, the division rate, $\sigma$ is set to be equal the natural death rate of the cells, $\mu$.

The state variables and parameters are summarised in Table S4 and in Figure 2a in the main text.

##### S1.1.2 Immunity

To capture viral clearance, we introduce the immune system through a single population of cytotoxic T-cells (CTLs) of size $T$. Their activity and dynamics are captured by the following ODE:

$$\dot{T} = \gamma\,(G+R)\,T\left(1-\frac{T}{K}\right) \tag{S2a}$$

CTLs grow at a rate of $\gamma$ in response to all infected cells. They also kill infected cells at a rate of $\alpha$ (see the infection equations below). We assume that the killing of cells primarily targets G and P stages, since cells spend little time in R stage relative to the other stages. To capture both clearance and immune escape by the virus, we assume that CTL growth is density dependant and include a carrying capacity, $K$.

##### S1.1.3 Small DNA viruses

To model small DNA oncoviruses, namely HPVs and the polyomaviruses, we assume the following model,

$$\dot{G} = \omega\,V + (2\,\delta_1 + \delta_2)\,R - (\sigma\,\varepsilon_2 + \mu\,(1-\varepsilon_1) + \alpha\,T)\,G \tag{S3a}$$

$$\dot{R} = \sigma\,\varepsilon_2\,G - (\delta_1 + \delta_2)\,R \tag{S3b}$$

$$\dot{P} = \delta_2\,R - (\mu + \alpha\,T)\,P \tag{S3c}$$

$$\dot{T} = \gamma\,(G+R+P)\,T\left(1-\frac{T}{K}\right) \tag{S3d}$$

$$\dot{V} = \phi\,P - \theta\,V. \tag{S3e}$$

Since small DNA viruses have two forms of resting infected cells, those with the virus in episomal form and those that produce virions, we include a new population of cells of size $P$ for the latter. We also track the total number of free viruses, $V$, which are produced by these cells. A schematic of this model is found in Figure 2b.

Resting cells become infected at a rate $\omega V$ because we assume that there is no depletion of the pool of susceptible cells (sDNA viruses infect tissues that have at least some regenerative capacity, such as the epithelium and kidneys [4]). The effects of viral oncogenes are represented by the two parameters. $\varepsilon_1$ scales down the natural death rate, $\mu$, thereby delaying or preventing cell death. The two mechanisms are oncogene anti-apoptotic programming and preventing innate immunity detection and killing. Thus, $\varepsilon_1$ captures two effects oncogenes have on the hallmarks of cancer [10]. The parameter $\varepsilon_2$ increases the rate of cell cycle re-entry (by increasing the natural rate, $\sigma$) and represents the viral oncogenes driving cell proliferation. Completion of cell division can lead to two outcomes. Either two cells at rest, $G$, are created at a rate $\delta_1$, or one cell at rest, $G$, and one productive cell, $P$, are created at rate $\delta_2$. We assume the immune system kills both $G$ and $P$ cells at an equal rate, $\alpha$, and that it grows in response to the presence of all classes of infected cells. Free virus is produced at a rate $\phi$ and cleared at a rate $\theta$.

The list of possible events, their rates of occurrence and their outcomes (associated updates in compartment sizes) for this model are shown in Table S1.

Table S1: **Small DNA virus model events and outcomes**

| **Rate** | **Event description** | **Outcome** |
|---|---|---|
| $\omega\ V$ | Infection of $G$ by free virus | $G \longrightarrow G+1$ |
| $2\ \delta_1\ R$ | Division results in 2 new $G$s | $G \longrightarrow G+2$<br>$R \longrightarrow R-1$ |
| $\delta_2\ R$ | Division results in 1$G$ and 1$P$ | $R \longrightarrow R-1$<br>$G \longrightarrow G+1$<br>$P \longrightarrow P+1$ |
| $\sigma\ \varepsilon_2\ G$ | $G$ moves into replication phase | $G \longrightarrow G-1$<br>$R \longrightarrow R+1$ |
| $\mu\ (1-\varepsilon_1)\ G$ | Natural death of $G$ | $G \longrightarrow G-1$ |
| $\alpha\ G\ T$ | Killing of $G$ by immunity | $G \longrightarrow G-1$ |
| $\mu\ P$ | Death of virus producing cells | $P \longrightarrow P-1$ |
| $\alpha\ P\ T$ | Killing of $P$ by immunity | $P \longrightarrow P-1$ |
| $\gamma\,(G+R+P)\ T\ (1-T/K)$ | Growth of $T$ cells from $G$ or $R$ or $P$ | $T \longrightarrow T+1$ |
| $\phi\ P$ | Production of free virions | $V \longrightarrow V+1$ |
| $\theta\ V$ | Clearance of virions | $V \longrightarrow V-1$ |

##### S1.1.4 Large DNA viruses

To model the infection dynamics of large DNA oncoviruses, such as KHSV and EBV, we use the following ODEs:

$$\dot{G} = \omega\, V + 2\, \delta\, R - (\sigma\, \varepsilon_2 + \mu\, (1-\varepsilon_1) + \eta + \alpha\, T)\, G \tag{S4a}$$

$$\dot{R} = \sigma\, \varepsilon_2\, G - \delta\, R \tag{S4b}$$

$$\dot{P} = \eta\, G - \alpha\, P\, T \tag{S4c}$$

$$\dot{T} = (\gamma_1\, (G+R) + \gamma_2\, P)\, T \left(1 - \frac{T}{K}\right) \tag{S4d}$$

$$\dot{V} = \phi\, P - \theta\, V. \tag{S4e}$$

The resting cells, $G$, of these viruses are in a latent state (the virus is again in episomal form) and the lytic phase only appears sporadically (i.e. we assume $\eta$ to be low and set it to 1/5 years$^{-1}$). Following biological evidence [10], we assume that the immune response is significantly more sensitive to productive cells (in lytic phase) than those in latent phase and therefore split $\gamma$ into two parameters, such that $\gamma = \gamma_1 \ll \gamma_2$. The schematic of this model is shown in Figure 2c in the main text.

The events of this model, their rates and outcomes are described in Table S2.

##### S1.1.5 Large DNA virus dynamics with memory

The classical Gillespie algorithms are memory-less, which can be problematic when calculating the waiting time of events such as maturation period that cannot occur very rapidly 'by chance'. Modifications of the algorithm allow to implement non-exponentially-distributed waiting times between events [5], unfortunately they do not apply to the $\tau$-leap algorithm, which we use here to avoid tracking cells individually.

To incorporate memory into the processes, we resort to a classical technique, which consists in splitting a compartment into sub-compartment (see [8] and Box 22.3 in [6]). By combining several exponential distributions (one to switch from each sub-compartment to the next), we converge towards a gamma distribution.

Table S2: **Large DNA virus model events and outcomes**

| Rate | Event description | Outcome |
|---|---|---|
| $\omega\, V$ | Infection of $G$ by free virus | $G \longrightarrow G+1$ |
| $2\, \delta\, R$ | Division results in 2 new $G$s | $G \longrightarrow G+2$<br>$R \longrightarrow R-1$ |
| $\sigma\, \varepsilon_2\, G$ | $G$ moves into replication phase | $G \longrightarrow G-1$<br>$R \longrightarrow R+1$ |
| $\mu\, (1-\varepsilon_1)\, G$ | Natural death of $G$ | $G \longrightarrow G-1$ |
| $\alpha\, G\, T$ | Killing of $G$ by immunity | $G \longrightarrow G-1$ |
| $\eta\, G$ | $G$ cell becoming producing cell | $P \longrightarrow P+1$<br>$G \longrightarrow G-1$ |
| $\alpha\, P\, T$ | Killing of $P$ by immunity | $P \longrightarrow P-1$ |
| $\gamma_1\, (G+R)\, T\, (1-T/K)$ | Growth of $T$ cells from $G$ or $R$ | $T \longrightarrow T+1$ |
| $\gamma_2\, P\, T\, (1-T/K)$ | Growth of $T$ cells from $P$ | $T \longrightarrow T+1$ |
| $\phi\, P$ | Production of free virions | $V \longrightarrow V+1$ |
| $\theta\, V$ | Clearance of virions | $V \longrightarrow V-1$ |

Mathematically, we introduce $s \in \mathbb{N}^*$ age classes for $G$ cells, each with a population size $G_s$. For $i > 1$, we move from $G_{i-1}$ to $G_i$ at a rate $\eta/s$, where is the rate at which latent cells ($G$) move into lytic phase ($P$) in the memory-less process. The final update is from $G_s$ to $P$ and it also occurs at a rate $\eta/s$. If $s = 1$, where are back to the memory-less process.

Notice that for updating purposes, the age class of the replicating cells also needs to be followed, which means introducing $s$ populations $R_s$.

##### S1.1.6 Retro-like viruses

We model the dynamics of retro-like oncoviruses, such as HTLV-1 and HBV, using the following set of ODEs:

$$\dot{G_p} = \omega\, V + 2\, \delta\, R - (\sigma\, \varepsilon_2 + \mu\, (1-\varepsilon_1) + \alpha\, T)\, G_p \tag{S5a}$$

$$\dot{R} = \sigma\, \varepsilon_2\, G_p - \delta\, R \tag{S5b}$$

$$\dot{T} = \gamma\, (G_p + R)\, T \left(1 - \frac{T}{K}\right) \tag{S5c}$$

$$\dot{V} = \phi\, G_p - \theta\, V. \tag{S5d}$$

Here, the infected resting cell population is given a subscript, $G_p$, because they bud newly formed virions. As before, infected resting cells can be produced by virions infecting new cells at a rate $\omega$. Cells enter replication at a rate $\varepsilon_2\, \sigma$ and die at a rate $\mu\, (1-\varepsilon_1)$ from natural death (and innate immunity killing) or at a rate $\alpha$ from the adaptive response, namely by CTLs. Replicating cells return to their resting state at a rate $\delta$ and produce two daughter cells. The total number of divisions is tracked using the equation $\dot{N} = \delta_1\, R$. See Figure 2d for the schematic of this model.

The events occurring in this model, their rates and corresponding outcomes are summarised in Table S3.

##### S1.1.7 Model parameter values

More generally, parameter values are taken to be biologically realistic. Most of these are discussed in a recent study [11]. All the variables and parameters are summarised in Table S4. We switch from one class of virus to another by adding or removing 2 parameters among a list of three ($\eta$, $\delta_2$ and $\gamma_2$) and for all of these we are able to make biologically-relevant assumptions regarding their default value.

Table S3: **Retro-like virus model events and outcomes**

| **Rate** | **Event description** | **Outcome** |
|---|---|---|
| $\omega\, V$ | Infection of $G_p$ by free virus | $G_p \longrightarrow G_p + 1$ |
| $2\, \delta\, R$ | Division results in 2 new $G_p$s | $G_p \longrightarrow G_p + 2$<br>R $\longrightarrow R - 1$ |
| $\sigma\, \varepsilon_2\, G_p$ | $G_p$ moves into replication phase | $G_p \longrightarrow G_p - 1$<br>R $\longrightarrow R + 1$ |
| $\mu\, (1-\varepsilon_1)\, G_p$ | Natural death of $G_p$ | $G_p \longrightarrow G_p - 1$ |
| $\alpha\, G_p\, T$ | Killing of $G_p$ by immunity | $G_p \longrightarrow G_p - 1$ |
| $\gamma\, (G_p + R)\, T\, (1 - T/K)$ | Growth of T cells from $G_p$ or $R$ | T $\longrightarrow T + 1$ |
| $\phi\, G_p$ | Production of free virions | V $\longrightarrow V + 1$ |
| $\theta\, V$ | Clearance of virions | V $\longrightarrow V - 1$ |

Table S4: **Variables and parameters of the models.** Values shown in the second column are default. We made an effort to homogenise these values as much as possible to compare life cycles. †0.5 for the retro-like viruses life cycle. * Besides natural cell death this rate also includes background killing by the innate immune response. ** Cell survival is due to viral shut down of apoptosis signalling and avoiding innate immunity destruction. *** This oncogene effect implies sustained proliferative signalling.

| **Variable** | | **Description** |
|---|---|---|
| $G$ & $G_U$ | | number of cells at rest, infected or uninfected with viral episomes |
| $G_p$ | | number of infected cells at rest that are budding virions |
| $R$ & $R_U$ | | number of cells undergoing cell division, infected or uninfected |
| $P$ | | number of productively infected cells (lytic or non-lytic) |
| $V$ | | number of free virions |
| $T$ | | number of cytotoxic T cells |
| $D$ | | number of replication rounds undergone by a cell |
| **Parameter** | **Default** | **Description** |
| $\sigma$ | $1/50$ | baseline rate at which cells at rest enter cell cycle for division (in days$^{-1}$) |
| $\mu$ | $1/50$ | baseline death rate of uninfected cells* (in days$^{-1}$) |
| $\delta$ | 1 | rate at which cell division produces two $G$ cells (in days$^{-1}$) |
| $\delta_1$ | 5/6 | same as $\delta$ for large-DNA |
| $\delta_2$ | $1/6$ | rate at which a cell division produces one $G$ cell and one $P$ cell (in days$^{-1}$) |
| $\eta$ | $1/1825$ | rate at which latent cells move into lytic phase (in days$^{-1}$) |
| $\gamma$ or $\gamma_1$ | $10^{-7}$ | CTL activation rate (in infected cell$^{-1}$.days$^{-1}$) |
| $\gamma_2$ | $10^{-6}$ | CTL activation rate by $P$ cells (in productive cell$^{-1}$.days$^{-1}$) |
| $\alpha$ | $10^{-3}$ | killing rate of infected cells by CTLs (in infected cell$^{-1}$.days$^{-1}$) |
| $K$ | 50 | CTL maximum population size (in cells) |
| $K_2$ | $10^{10}$ | Infected cell maximum population size (in cells) |
| $\phi$ | 0.5 or 2† | Virion producing rate by $P$ cells (in days$^{-1}$) |
| $\theta$ | 1/2 | Virion clearance rate (in days$^{-1}$) |
| $\omega$ | $10^{-3}$ | Infection rate (in days$^{-1}$) |
| $\varepsilon_1$ | $\in [0,1]$ | oncogene effect acting against cell death** |
| $\varepsilon_2$ | $\in [1,5]$ | oncogene effect acting on entrance into cell cycle and cell proliferation*** |
| $\nu$ | $5 \cdot 10^{-15}$ | scaling parameter for cancer risk (in $G$ cell$^{-1}$.days$^{-1}$) |
| $p$ | 1 | exponent capturing the relationship between $D$ and cancer risk |

#### S1.2 Between-host modelling and numerical implementation

##### S1.2.1 Cancer initiation events

We explicitly include a 'catastrophic' event corresponding to a cancer initiation event. For simplicity, instead of explicitly tracking the accumulation of mutations that drive cells towards a cancerous state [2], we track the number of the replication cycles an infected cell has gone through. We then assume that, at a given time $t$, each infected cell can trigger a cancer at a rate $\nu D^p$, where $\nu = 5 \cdot 10^{-15}$ is a scaling parameter, $D$ is the number of divisions undergone by the cell and $p$ is a scaling parameter between $D$ and cancer risk. This assumption is motivated by results showing that the lifetime risk of cancer in a tissue correlated with the number of stem cell divisions in the lifetime of the tissue [12]. In the main text, we assume that $p = 1$ (linear correlation) but in Supplementary Results we investigate an absence of correlation ($p = 0$) or non-linear relationships ($p = 2$ and $p = 3$). In practice, in the Gillespie algorithm, the $G$ and $R$ populations are split into subpopulations ($G_D$ and $R_D$) such that when a cell enters division we have a transition from $G_D$ to $R_D$ and when a division ends we produce two cells, one in $G_D$ and another in $G_{D+1}$.

As indicated in the main text, a possible extension of this model, in addition to adding several steps to reach the cancer state, would be to explicitly follow the growth of a population of cancer cells, $C$, generated through this process.

###### S1.2.2 Between-host transmission event

In order to go beyond the within-host level, we include a transmission process. We assume that between-host contacts that are sufficiently close for transmission take place regularly (every $\Delta t = 30$ days). The probability of transmission, $p$, at each contact is given by $p(t) = V(t)/(10^4 + V(t))$. This type II functional response assumption means that transmission probability first increases linearly with virus load but then reaches a plateau for high virus loads. In the case of HPV, this plateau is likely to be reached rapidly since the probability of transmission per sex act is estimated to be close to 90% [13].

We also investigated less frequent transmission ($\Delta t = 60$ days), which led to similar results. Conversely, increasing transmission opportunities leads to an alignment between within-host and between-host fitness (Figure not shown).

###### S1.2.3 Gillespie algorithm

Given the number of cells involved, we use the $\tau$-leap Gillespie algorithm based on the events, rates and outcomes listed in Table S1, S2 and S3. We use a fixed time step of 0.5 day. Simulations are run until viral clearance, cancer initiation event, or the time limit (set to 50 years) is reached.

The discrete and numerical implementation allows us to readily follow the number of division each cell has gone through. This means tracking a vector of size $D$, where $D$ is the number of divisions the cell has been through, for $G$ and $R$ cells. We are therefore working with subpopulations ($G_D$ and $R_D$). For low populations sizes, the update is performed stochastically: we first determine how many replication events are happening at time $t$ and then we select which of the many cells in the $D$ subpopulations are involved based on their relative frequency. When the population size of $G$ cells becomes greater than $10^4$, we assign the replication events deterministically but still based on the relative frequency of the subpopulations.

We explore 984 parameter sets with different values of $\varepsilon_1 \in [0,1]$ and $\varepsilon_2 \in [0.5,5]$, assuming uniform distributions for each. Notice that in the main text we do not report situations where the oncovirus decreases the proliferation rate of the infected cell ($\varepsilon_2 < 1$). Each parameter set was simulated 25 times.

#### S2 Supplementary Results

##### S2.1 Number of cell divisions and cancer

In Figure S1, we present the stable distribution of the number of divisions the cells have been through in the population (i.e. the $G_D$ subpopulations). As expected, this distribution is log-linear. We also see that with our default parameter values, the distributions level off before reaching 30 divisions, which is considered to be an upper limit for many cells (see figure caption). This threshold is reached because for large total populations of cells ($> 10^4$) our updating it not stochastic, such that small cell populations ($< 20 cells$) become negligible (in the figure there are more than 5 orders of magnitude difference between the most abundent population and the rare ones). Notice that we assume cells having divided 30 times or more do not produce any offspring anymore.

We also see that the action of the oncogenes to increase cell life expectancy ($\varepsilon_1$) shifts this distribution to the right. Even a maximum efficiency of $\varepsilon_1$ does not make cells truly immortal since they are still targeted by the adaptive immune response.

We then vary the effect of the number of divisions undergone by a cell on the risk that it becomes cancerous. In the main text, we assumed a linear relationship ($p = 1$), based on empirical data [12]. Here, we also study

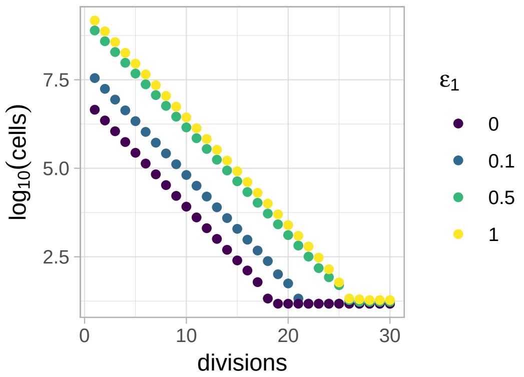

Figure S1: **Distribution of the number of divisions undergone by infected cells.** $\varepsilon_1$ increases this number of divisions by decreasing cell mortality rate. Notice that the distributions level off before 30 divisions, which is the maximum threshold.

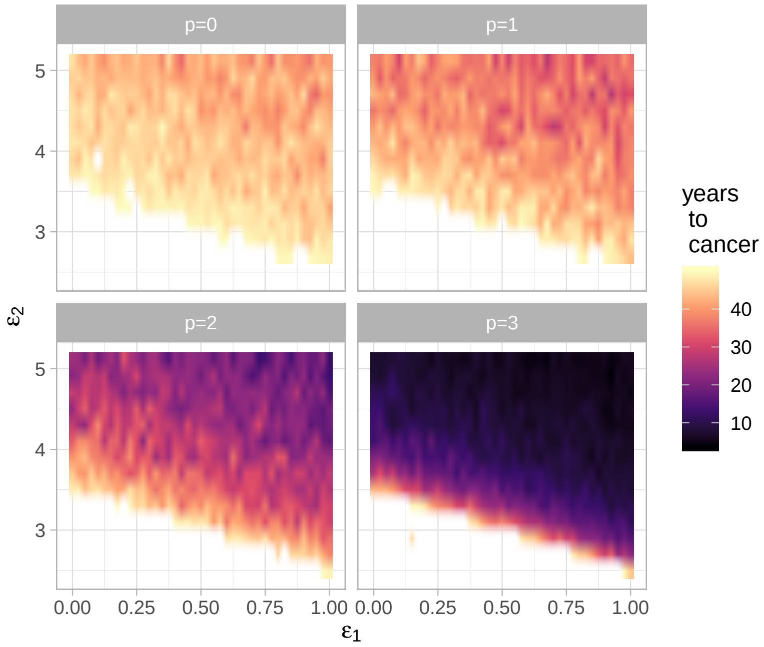

Figure S2: **Link between number of cell divisions and cancer risk.** We vary the exponent $p$, which is related to cancer risk with the formula $\nu \sum_D G_D D^p$, where $\nu$ is a constant scaling parameter, $D$ is the number of divisions undergone by cells from the $G_D$ population.

an absence of correlation (i.e. the number of divisions has no effect on cancer risk, $p = 0$) and non-linear relationships ($p > 1$). As shown in Figure S2, the value of $p$ does not seem to affect the probability that cancer will occur given a parameter set $(\varepsilon_1, \varepsilon_2)$. This is somehow surprising because we could have expected to see an interaction between $p$ and oncogene action. However, $p$ has a dramatic effect on the speed at which the cancer initiation event occurs.

This limited qualitative effect of $p$ on the result suggests that the action of $\varepsilon_1$ to increase infected cell life expectancy is still too limited to significantly alter the distribution of the number of divisions (see also Figure S1). Considering a third action of the oncogenes, namely evading adaptive immunity, would most likely make the exact link between cell divisions and cancer much more more important.

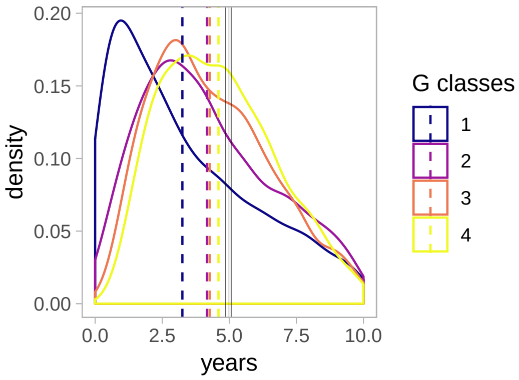

Figure S3: **For large-DNA viruses, distribution of the waiting time for a *G* cell to become a *P* cell.** As the number of intermediate steps ($s$) to switch from $G$ to $P$ increases, the distributions change and the median (dashed line) becomes closer to the mean (full line).

#### S2.2 Effect of memory processes for large-DNA viruses

As explained above, we add sub-populations to the $G$ and $R$ population in order to track the age of the cell and to introduce memory into the Gillespie algorithm. In practice, $G$ cells have to go through $s$ different classes before being able to produce virions.

Figure S3 shows that the distribution of the time until a $P$ cell is produced from an initial $G$ cell is greatly affected by the number of classes. For a single class ($s = 1$, our default value), the median of the distribution (dashed line) is far from the mean (full line). Biologically, this means that this event can occur very rapidly (in less than a year) by chance. Increasing the number of classes increases the median value towards mean. We also see that the event almost never happens until a year by chance.

The number of classes for $G$ (that is the value of $s$) has a strong effect on the selective pressures acting on the virus. Intuitively, since virions cannot be produced rapidly at all, any strategy that decreases infection duration is extremely strongly counter-selected. Furthermore, being able to increase the average age of the infected cell population, for instance by decreasing apoptosis, should be favoured.

To illustrate this, we ran simulations for our 984 parameter combinations for 4 different values of $s$. We then selected the 50 combinations that achieved the highest between-host fitness (independently of whether cancer occurred or not). In Figure S4a, we see that delaying the median time until virion production strongly decreases fitness. We then investigate the viral strategies in these fittest combinations. Recall that in the main text, prolonging the life expectancy of the cell bore no clear signal for large-DNA viruses. Here, we see that increasing $s$ leads to higher virus manipulation of cell mortality (Figure S4b). With only 4 classes, most of the $\varepsilon_1$ value are close to the maximum. Increasing the number of $G$ classes also increases the value of $\varepsilon_2$ in the fittest parameter sets, although only for $s \geq 3$ (Figure S4c). This could be due to the fact that $\varepsilon_1$ is not sufficient to compensate for the constraint imposed by the late production of virions.

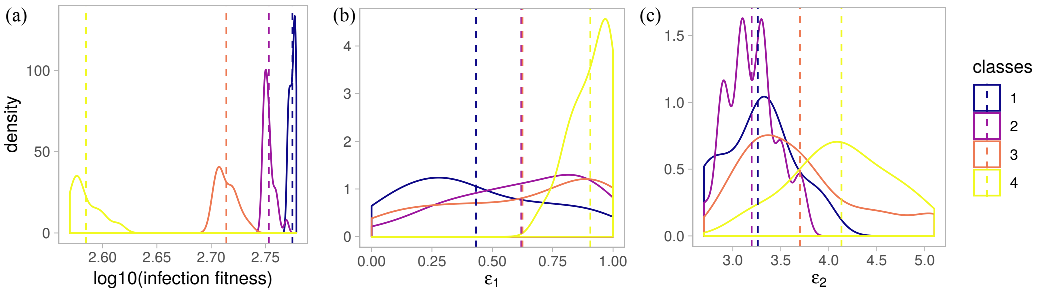

Figure S4: **For large-DNA viruses, effect of the number of $G$ subpopulation on a) infection fitness, b) $\varepsilon_1$ and c) $\varepsilon_2$.** these distributions are calculated on the 50 fittest parameter sets. Dashed lines show mean values.
