## Supplementary material for "Modelling the evolution of viral oncogenesis": R scripts to resimulate the data and generate the plots: get_sum_stats_retrovir.R

```
library(ggplot2)

setwd("/home/samuel/Documents/Hote-parasite/HPV/PTRS/Evolution_oncoviruses/Analysis")


# data1<-read.csv(file="allruns_lDNA_1classes.csv")
# data2<-read.csv(file="allruns_lDNA_2classes.csv")
# data3<-read.csv(file="allruns_lDNA_4classes.csv")
# data4<-read.csv(file="allruns_lDNA_10classes.csv")
# data_raw<-rbind(data1,data2,data3,data4)

#write.csv(data_raw,file="allruns_lDNA_new.csv",row.names=F)


#data_raw<-read.csv(file="allruns_retrovir.csv")
data_raw<-read.csv(file="output_retrovir_div2.csv")


data<-as.data.frame(data_raw)

data$cancer<-as.numeric(data$mut>0)

head(data)


names(data)
library(ggplot2)

#duration for all values of epsilon
ggplot(data, aes(log10(tfinal)))+
  scale_colour_viridis_d()+
  geom_density()


ggplot(data, aes(log10(trans+1)))+
  scale_colour_viridis_d()+
  geom_density()

ggplot(data, aes((mut>0)))+
  scale_colour_viridis_d()+
  geom_density()


#creating average summary statistics
sumstats<-c()
sumstats_median<-c()


    for(i1 in unique(data$epsilon1))
    {
      for(i2 in unique(data$epsilon2))
      {
        temp<-subset(data,(epsilon1==i1)&(epsilon2==i2))
        
        if(dim(temp)[1]>0)
        {
          sumstats<-rbind(sumstats,colSums(temp)[-1]/dim(temp)[1])
          
          sumstats_median<-rbind(sumstats_median,apply(temp,2,FUN=median)[-1])
        }
        
      }
    }


sumstats<-as.data.frame(sumstats)
sumstats_median<-as.data.frame(sumstats_median)

write.csv(sumstats,file="sumstats_mean_retrovir2.csv",row.names = FALSE)
write.csv(sumstats_median,file="sumstats_median_retrovir2.csv",row.names = FALSE)


#############
#Plot time to cancer
#############

#for all the number of classes (for comparison)     
ggplot(subset(sumstats_median,as.numeric(mut)>0), 
       aes(epsilon1,epsilon2,z=tfinal))+
  geom_raster(aes(fill = tfinal)) +
  #  geom_contour(colour = "white")+
  labs(fill="time to cancer",x=expression(paste(epsilon [1])),y=expression(paste(epsilon [2])))+
  theme(axis.text=element_text(size=10),axis.title=element_text(size=14),plot.title=element_text(size=12,face="bold"))


ggplot(subset(sumstats_median,(mut>0)), 
       aes(epsilon1,epsilon2,z=tfinal/365))+
  geom_raster(aes(fill = tfinal/365)) +
  #  geom_contour(colour = "white")+
  labs(fill="years to cancer",x=expression(paste(epsilon [1])),y=expression(paste(epsilon [2])))+
  theme(axis.text=element_text(size=10),axis.title=element_text(size=14),plot.title=element_text(size=12,face="bold"))

#ggsave(plot=last_plot(),filename="Time_to_cancer_N1.pdf", width = 11, height = 8, units = "cm")
```
