## Supplementary material for "Modelling the evolution of viral oncogenesis": R scripts to resimulate the data and generate the plots: plot_single_run_allviruses_clearance.R

```
setwd("/home/samuel/Documents/Hote-parasite/HPV/PTRS/Evolution_oncoviruses/Gillespie/single_runs")


##########################
#load libraries
##########################
library(ggplot2)


gdata1<-read.csv(file = "single_run_lDNA_1_cleared_(05-1).csv",header=T)
gdata2<-read.csv(file = "single_run_sDNA_cleared_(05-1).csv",header=T)
gdata3<-read.csv(file = "single_run_retrovir_cleared_(05-1).csv",header=T)

gdata1$virus<-"large-DNA"
gdata2$virus<-"small-DNA"
gdata3$virus<-"retro-like"


gdata<-rbind(gdata1,gdata2,gdata3)
gdata<-as.data.frame(gdata)

allruns<-unique(gdata$run)
iruns<-max(allruns)


Tmax <- 18250 
numGclasses<-as.integer(unique(gdata$N))


alldata<-gdata

#keep only keely counts
small_data<-gdata[round(gdata$time)==(gdata$time),]
small_data<-as.data.frame(small_data)

names(small_data)

#ggplot(gdata,
ggplot(subset(small_data,(as.numeric(run)==4)),
       aes(as.numeric(as.character(time))/365,as.numeric(as.character(popsize)),colour=as.factor(type),linetype=as.factor(run))
)+
  geom_line(size=0.2)+
  guides(linetype=FALSE)+
  labs(colour="cells",x="time (years)",y=expression(paste(log[10]("population size+1"))),linetype="run")+
  theme(axis.text=element_text(size=10),axis.title=element_text(size=14),plot.title=element_text(size=12,face="bold"))+
  scale_color_viridis_d()+
  facet_wrap(~as.factor(virus))

#ggplot(gdata,
ggplot(small_data,
              aes(as.numeric(as.character(time))/365,as.numeric(as.character(popsize)),colour=as.factor(type),linetype=as.factor(run))
)+
  geom_line(size=0.2)+
  guides(linetype=FALSE)+
  labs(colour="cells",x="time (years)",y=expression(paste(log[10]("population size+1"))),linetype="run")+
  theme(axis.text=element_text(size=10),axis.title=element_text(size=14),plot.title=element_text(size=12,face="bold"))+
  scale_color_viridis_d(option="inferno")+
  theme_light()+
  facet_wrap(~as.factor(virus),scale="free")


ggplot(subset(small_data,(as.numeric(run)==c(1:3))),
       aes(as.numeric(as.character(time)),as.numeric(as.character(popsize)),colour=as.factor(type),linetype=as.factor(run))
)+
  geom_line(size=0.3)+
  #  ylim(1,10)+
  labs(colour="cells",x="days",y=expression(paste(log[10]("population size+1"))),linetype="run")+
  scale_color_viridis_d(option="plasma")+
  theme_light()+
  theme(panel.grid.major = element_blank(),panel.grid.minor = element_blank(),legend.position = "none")+
  facet_wrap(~as.factor(virus),scale="free")


 ggsave(plot=last_plot(),filename="single_run_1_cleared.pdf", width = 14.0, height = 5.0, units = "cm")


 
 ggplot(subset(small_data,(as.numeric(run)==c(1:3))),
        aes(as.numeric(as.character(time)),10^as.numeric(as.character(popsize)),colour=as.factor(type),linetype=as.factor(run))
 )+
   geom_line(size=0.3)+
   scale_y_log10(   breaks = c(10, 1000, 100000), labels = expression(10, 10^3, 10^5))+
   labs(colour="cells",x="days",y="population size",linetype="run")+
   scale_color_viridis_d(option="plasma")+
   theme_light()+
   theme(panel.grid.major = element_blank(),panel.grid.minor = element_blank(),legend.position = "none")+
   facet_wrap(~as.factor(virus),scale="free")
 
 ggsave(plot=last_plot(),filename="single_run_1_cleared.pdf", width = 14.0, height = 5.0, units = "cm")
 


 ggplot(small_data,
        aes(as.numeric(as.character(time))/365,as.numeric(as.character(popsize)),colour=as.factor(type))
 )+
   geom_smooth()+
   guides(linetype=FALSE)+
   labs(colour="cells",x="time (years)",y=expression(paste(log[10]("population size+1"))),linetype="run")+
   theme(axis.text=element_text(size=10),axis.title=element_text(size=14),plot.title=element_text(size=12,face="bold"))+
   scale_color_viridis_d(option="inferno")+
   #  geom_vline(mapping=NULL,data=vdata,xintercept=as.vector(vdata$time/365),aes(linetype=as.factor(vdata$run)),size=0.5,colour=5)+
   theme_light()+
   facet_wrap(~as.factor(virus),scale="free")
 
 
 #ggsave(plot=last_plot(),filename="single_run_sDNA_cleared_fit.pdf", width = 11, height = 8, units = "cm")
 
 ggsave(plot=last_plot(),filename="single_run_cleared_fit.pdf", width = 17, height = 6.5, units = "cm")
 
library(data.table)
names(gdata)
temp=dcast(gdata,time+N+run~type,value.var="popsize")

head(temp)
head(gdata)


pdf()

plot(rep(-10,length(c(0:round(Tmax/365))))~c(0:round(Tmax/365)),type="l",xlab="time (years)",ylab="log[10](population size +1)",xlim=c(0,round(Tmax/365)),ylim=c(0,9))
for(j in c(1:7))
{
  data<-as.data.frame(results[[j]])
  timeY<-as.numeric((data$time/365))
  #print(max(timeY))
  lines(log10(1+data$G)~timeY,col=1,lwd=0.2)
  lines(log10(1+data$R)~timeY,col=2,lwd=0.2)
  lines(log10(1+data$V)~timeY,col=3,lwd=0.2)
  lines(log10(1+data$Tc)~timeY,col=4,lwd=0.2)
 # abline(v=max(timeY),lty=2)
}

dev.off()
```
