## Supplementary material for "Modelling the evolution of viral oncogenesis": R scripts to resimulate the data and generate the plots: plot_single_runs_allviruses_homoestasis.R

```
setwd("/home/samuel/Documents/Hote-parasite/HPV/PTRS/Evolution_oncoviruses/Gillespie/single_runs")


##########################
#load libraries
##########################
library(ggplot2)


gdata1<-read.csv(file = "single_run_lDNA_1_cleared_(0-1).csv",header=T)
gdata2<-read.csv(file = "single_run_sDNA_cleared_(0-1).csv",header=T)
gdata3<-read.csv(file = "single_run_retrovir_cleared_(0-1).csv",header=T)

gdata1$virus<-"large-DNA"
gdata2$virus<-"small-DNA"
gdata3$virus<-"retro-like"


gdata<-rbind(gdata1,gdata2,gdata3)
gdata<-as.data.frame(gdata)

allruns<-unique(gdata$run)
iruns<-max(allruns)


Tmax <- 18250 

# vdata<-c()
# for(ig in allruns)
# {
#   timeY<-max(gdata$time[gdata$run==ig])
#   if(max(timeY)<Tmax)
#   {
#     vdata<-rbind(vdata,c(max(timeY),numGclasses,ig))
#   }
# }
# 
# colnames(vdata)<-c("time","N","run")
# rownames(vdata)<-c()
# vdata<-as.data.frame(vdata)


names(gdata)

alldata<-gdata

#keep only keely counts
small_data<-gdata[round(gdata$time/7)==(gdata$time/7),]
small_data<-as.data.frame(small_data)


#keep a weekly resolution for the first 500 days
small_data1<-small_data[small_data$time<=1000,]

#then a 4 weeks resolution
small_data2<-small_data[(small_data$time>1000)&(small_data$time<=365*30),]
small_data2<-small_data2[round(small_data2$time/28)==(small_data2$time/28),]

small_data3<-rbind(small_data1,small_data2)
small_data3<-as.data.frame(small_data3)


#ggplot(gdata,
ggplot(subset(small_data3,(as.numeric(run)==c(1:5))),
       aes(log10(as.numeric(as.character(time))),as.numeric(as.character(popsize)),colour=as.factor(type),linetype=as.factor(run))
)+
  geom_line(size=0.2)+
  guides(linetype=FALSE)+
  labs(colour="cells",x="time (years)",y=expression(paste(log[10]("population size+1"))),linetype="run")+
  theme(axis.text=element_text(size=10),axis.title=element_text(size=14),plot.title=element_text(size=12,face="bold"))+
  scale_color_viridis_d()+
  facet_wrap(~as.factor(virus))+
  theme_light()

#ggplot(gdata,
ggplot(subset(small_data3,(as.numeric(run)==c(1:3))),
       aes(log10(as.numeric(as.character(time))),as.numeric(as.character(popsize)),colour=as.factor(type),linetype=as.factor(run))
)+
  geom_line(size=0.3)+
#  ylim(1,10)+
#  guides(linetype=FALSE)+
  labs(colour="cells",x=expression(paste(log[10]("days"))),y=expression(paste(log[10]("population size+1"))),linetype="run")+
#  theme(axis.text=element_text(size=10),axis.title=element_text(size=14),plot.title=element_text(size=12,face="bold"))+
  scale_color_viridis_d(option="plasma")+
  theme_light()+
  theme(panel.grid.major = element_blank(),panel.grid.minor = element_blank(),legend.position = "none")+
  facet_wrap(~as.factor(virus),scale="free")


#ggplot(gdata,
ggplot(subset(small_data3,(as.numeric(run)==c(1:3))),
       aes(as.numeric(as.character(time)),10^as.numeric(as.character(popsize)),colour=as.factor(type),linetype=as.factor(run))
)+
  geom_line(size=0.3)+
  #  ylim(1,10)+
  #  guides(linetype=FALSE)+
  scale_y_log10(   breaks = c(1,10, 10^2, 10^3, 10^4, 10^5), labels = expression(1,10,10^2, 10^3, 10^4, 10^5))+
  scale_x_log10(   breaks = c(1, 10, 100, 1000, 10000), labels = expression(1, 10, 10^2, 10^3, 10^4))+
  labs(colour="cells",x="days",y="population size",linetype="run")+
  #  theme(axis.text=element_text(size=10),axis.title=element_text(size=14),plot.title=element_text(size=12,face="bold"))+
  scale_color_viridis_d(option="plasma")+
  theme_light()+
  theme(panel.grid.major = element_blank(),panel.grid.minor = element_blank(),legend.position = "none")+
  facet_wrap(~as.factor(virus))


ggsave(plot=last_plot(),filename="single_run_1_cleared_01.pdf", width = 14, height = 5, units = "cm")


ggplot(subset(small_data3,(as.numeric(run)!=9)),
       aes(as.numeric(as.character(time))/365,as.numeric(as.character(popsize)),colour=as.factor(type))
)+
  geom_smooth()+
  guides(linetype=FALSE)+
  labs(colour="cells",x="time (years)",y=expression(paste(log[10]("population size+1"))),linetype="run")+
  theme(axis.text=element_text(size=10),axis.title=element_text(size=14),plot.title=element_text(size=12,face="bold"))+
  scale_color_viridis_d(option="inferno")+
  #  geom_vline(mapping=NULL,data=vdata,xintercept=as.vector(vdata$time/365),aes(linetype=as.factor(vdata$run)),size=0.5,colour=5)+
  theme_light()+
  facet_wrap(~as.factor(virus),scale="free")


#ggsave(plot=last_plot(),filename="single_run_sDNA_cleared_fit.pdf", width = 11, height = 8, units = "cm")

ggsave(plot=last_plot(),filename="single_run_fit.pdf", width = 17, height = 6.5, units = "cm")
```
