## Supplementary material for "Modelling the evolution of viral oncogenesis": R scripts to resimulate the data and generate the plots: plotting_lDNA_classes_effect.R

```
library(grid)
library(gridExtra)
library(ggplot2)

#set your working directory
setwd("/home/samuel/Documents/Hote-parasite/HPV/PTRS/Evolution_oncoviruses/Analysis")


#load files
sumstats<-read.csv(file="sumstats_mean_lDNA1_1.csv",header = TRUE)
sumstats_median<-read.csv(file="sumstats_median_lDNA1_1.csv",header = TRUE)
sumstats$rel_trans<-sumstats$trans/max(sumstats$trans)
sumstats_median$rel_trans<-sumstats_median$trans/max(sumstats_median$trans)

sumstats2<-read.csv(file="sumstats_mean_lDNA2_1.csv",header = TRUE)
sumstats_median2<-read.csv(file="sumstats_median_lDNA2_1.csv",header = TRUE)
sumstats2$rel_trans<-sumstats2$trans/max(sumstats2$trans)
sumstats_median2$rel_trans<-sumstats_median2$trans/max(sumstats_median2$trans)

sumstats3<-read.csv(file="sumstats_mean_lDNA3_1.csv",header = TRUE)
sumstats_median3<-read.csv(file="sumstats_median_lDNA3_1.csv",header = TRUE)
sumstats3$rel_trans<-sumstats3$trans/max(sumstats3$trans)
sumstats_median3$rel_trans<-sumstats_median3$trans/max(sumstats_median3$trans)


sumstats4<-read.csv(file="sumstats_mean_lDNA4_1.csv",header = TRUE)
sumstats_median4<-read.csv(file="sumstats_median_lDNA4_1.csv",header = TRUE)
sumstats4$rel_trans<-sumstats4$trans/max(sumstats4$trans)
sumstats_median4$rel_trans<-sumstats_median4$trans/max(sumstats_median4$trans)

sumstats8<-read.csv(file="sumstats_mean_lDNA8_1.csv",header = TRUE)
sumstats_median8<-read.csv(file="sumstats_median_lDNA8_1.csv",header = TRUE)
sumstats8$rel_trans<-sumstats8$trans/max(sumstats8$trans)
sumstats_median8$rel_trans<-sumstats_median8$trans/max(sumstats_median8$trans)


sumstats<-rbind(sumstats,sumstats2,sumstats3,sumstats4,sumstats8)
sumstats_median<-rbind(sumstats_median,sumstats_median2,sumstats_median3,sumstats_median4,sumstats_median8)


head(sumstats3)
head(sumstats4)

sumstats<-as.data.frame(sumstats)
sumstats_median<-as.data.frame(sumstats_median)


#############
#Plot Log(infection duration)
#############

  #for all the number of classes (for comparison)     
  ggplot(sumstats_median, 
       aes(as.numeric(epsilon1),as.numeric(epsilon2),z=as.numeric(log10(tfinal))))+
  geom_raster(aes(fill = as.numeric(log10(tfinal))), interpolate = FALSE) +
#  geom_contour(colour = "white",binwidth = .2)+
  labs(fill=expression(paste(log[10] (T[final]) )),x=expression(paste(epsilon [1])),y=expression(paste(epsilon [2])))+
  theme(axis.text=element_text(size=10),axis.title=element_text(size=14),plot.title=element_text(size=12,face="bold"))+
  facet_wrap(~ as.factor(numGclasses))+
  scale_fill_viridis_c(option="inferno")+
  theme_light()

ggsave(plot=last_plot(),filename="T_final_lDNA_all.pdf", width = 18, height = 10, units = "cm")


  #no effect of N on infection duration patterns

#for one number of G classes
ggplot(subset(sumstats_median,(as.factor(numGclasses)==4)), 
       aes(as.numeric(epsilon1),as.numeric(epsilon2),z=as.numeric(log10(tfinal))))+
  geom_raster(aes(fill = as.numeric(log10(tfinal))), interpolate = FALSE) +
#  geom_contour(colour = "white",binwidth = .2)+
  labs(fill=expression(paste(log[10] (T[final]) )),x=expression(paste(epsilon [1])),y=expression(paste(epsilon [2])))+
  theme(axis.text=element_text(size=10),axis.title=element_text(size=14),plot.title=element_text(size=12,face="bold"))+
  scale_fill_viridis_c(option="inferno")+
  theme_minimal()

#ggsave(plot=last_plot(),filename="T_final_lDNA_N4_div3.pdf", width = 11, height = 8, units = "cm")


#############
#Plot BH fitness
#############

#for all the number of classes (for comparison)     
ggplot(sumstats_median, 
       aes(epsilon1,epsilon2,z=log10(trans+1)))+
  geom_raster(aes(fill = log10(trans+1)), interpolate = FALSE) +
  #geom_contour(colour = "white")+
  labs(fill="fitness",x=expression(paste(epsilon [1])),y=expression(paste(epsilon [2])))+
facet_wrap(~ as.factor(numGclasses))+
  scale_fill_viridis_c(option="plasma")+
  theme_light()

ggsave(plot=last_plot(),filename="BH_Fitness_lDNA_all.pdf", width = 18, height = 10, units = "cm")


#for all the number of classes (for comparison)     
ggplot(sumstats_median, 
       aes(epsilon1,epsilon2,z=rel_trans))+
  geom_raster(aes(fill = rel_trans), interpolate = FALSE) +
  #geom_contour(colour = "white")+
  labs(fill="% max trans",x=expression(paste(epsilon [1])),y=expression(paste(epsilon [2])))+
  theme(axis.text=element_text(size=10),axis.title=element_text(size=14),plot.title=element_text(size=12,face="bold"))+
  facet_wrap(~ as.factor(numGclasses))+
  scale_fill_viridis_c(option="plasma")+
  theme_light()

#ggsave(plot=last_plot(),filename="relative_BH_Fitness_lDNA_all_div1.pdf", width = 17, height = 6.5, units = "cm")


# N strongly decreases BH fitness (less viruses being produced)


ggplot(subset(sumstats_median,(as.factor(numGclasses)==4)), 
       aes(epsilon1,epsilon2,z=trans))+
  geom_raster(aes(fill = trans),interpolate=TRUE) +
#  geom_contour(colour = "white",binwidth = 100)+
  labs(fill=expression(paste(Infections)),x=expression(paste(epsilon [1])),y=expression(paste(epsilon [2])))+
  scale_fill_viridis_c(option="plasma")+
  theme_minimal()

#ggsave(plot=last_plot(),filename="BH_Fitness_lDNA_N4_div3.pdf", width = 11, height = 8, units = "cm")


#############
#Plot WH fitness
#############

#for all the number of classes (for comparison)     
ggplot(sumstats_median, 
       aes(epsilon1,epsilon2,z=log10(Vtot+1)))+
  geom_raster(aes(fill = log10(Vtot+1)),interpolate = FALSE) +
  #  geom_contour(colour = "white")+
  labs(fill=expression(paste(log[10] (V[tot]) )),x=expression(paste(epsilon [1])),y=expression(paste(epsilon [2])))+
  facet_wrap(~ as.factor(numGclasses))+
  scale_fill_viridis_c(option="viridis")+
  theme_light()

ggsave(plot=last_plot(),filename="WH_Fitness_lDNA_all.pdf", width = 18, height = 10, units = "cm")


# N decreases WH fitness (less reinfection)


ggplot(subset(sumstats_median,(as.factor(numGclasses)==4)), 
       aes(epsilon1,epsilon2,z=log10(Vtot+1)))+
  geom_raster(aes(fill = log10(Vtot+1)),interpolate=TRUE) +
#  geom_contour(colour = "white")+
  labs(fill=expression(paste(log[10] (V[tot]) )),x=expression(paste(epsilon [1])),y=expression(paste(epsilon [2])))+
  theme(axis.text=element_text(size=10),axis.title=element_text(size=14),plot.title=element_text(size=12,face="bold"))+
  scale_fill_viridis_c(option="viridis")+
  theme_minimal()

#ggsave(plot=last_plot(),filename="WH_Fitness_lDNA_N4_div3.pdf", width = 11, height = 8, units = "cm")


#############
#Plot proportion cancer
#############


#for all the number of classes (for comparison)     
ggplot(sumstats, 
       aes(epsilon1,epsilon2,z=cancer))+
  geom_raster(aes(fill = cancer),interpolate=TRUE) +
#   geom_contour(colour = "white")+
  labs(fill=expression(paste(cancer)),x=expression(paste(epsilon [1])),y=expression(paste(epsilon [2])))+
  facet_wrap(~ as.factor(numGclasses))+
  scale_fill_viridis_c(option="cividis")+
  theme_light()

ggsave(plot=last_plot(),filename="Cancer_lDNA_all.pdf", width = 18, height = 10, units = "cm")


#no effect on cancer


ggplot(subset(sumstats,(as.factor(numGclasses)==4)), 
       aes(epsilon1,epsilon2,z=cancer))+
  geom_raster(aes(fill = cancer),interpolate=TRUE) +
#  geom_contour(colour = "white")+
  labs(fill=expression(paste(cancer)),x=expression(paste(epsilon [1])),y=expression(paste(epsilon [2])))+
  theme(axis.text=element_text(size=10),axis.title=element_text(size=14),plot.title=element_text(size=12,face="bold"))+
  scale_fill_viridis_c(option="cividis")+
  theme_minimal()

#ggsave(plot=last_plot(),filename="Cancer_lDNA_N4_div3.pdf", width = 11, height = 8, units = "cm")


#############
#Plot time to cancer
#############


#for all the number of classes (for comparison)     
ggplot(subset(sumstats,as.numeric(mut)>0), 
       aes(epsilon1,epsilon2,z=tfinal/365))+
  geom_raster(aes(fill = tfinal/365),interpolate=TRUE) +
  #  geom_contour(colour = "white")+
  labs(fill="years\n to\n cancer",x=expression(paste(epsilon [1])),y=expression(paste(epsilon [2])))+
  facet_wrap(~ as.factor(numGclasses))+
  scale_fill_viridis_c(option="magma")+
  theme_light()

ggsave(plot=last_plot(),filename="Time_to_cancer_lDNA.pdf", width = 18, height = 10, units = "cm")


#no effect of N on time to cancer


ggplot(subset(sumstats_median,(as.factor(numGclasses)==4)&(mut>0)), 
       aes(epsilon1,epsilon2,z=tfinal/365))+
  geom_raster(aes(fill = tfinal/365),interpolate=TRUE) +
  #  geom_contour(colour = "white")+
  labs(fill="years to cancer",x=expression(paste(epsilon [1])),y=expression(paste(epsilon [2])))+
  theme(axis.text=element_text(size=10),axis.title=element_text(size=14),plot.title=element_text(size=12,face="bold"))+
  scale_fill_viridis_c(option="magma")


#ggsave(plot=last_plot(),filename="Time_to_cancer_lDNA4_3.pdf", width = 11, height = 8, units = "cm")


#############
#Plot time to recovery
#############

#for all the number of classes (for comparison)     
ggplot(subset(sumstats_median,as.numeric(mut)==0), 
       aes(epsilon1,epsilon2,z=tfinal))+
  geom_raster(aes(fill = tfinal),interpolate=TRUE) +
  #  geom_contour(colour = "white")+
  labs(fill="time to recovery",x=expression(paste(epsilon [1])),y=expression(paste(epsilon [2])))+
  theme(axis.text=element_text(size=10),axis.title=element_text(size=14),plot.title=element_text(size=12,face="bold"))+
  facet_wrap(~ as.factor(numGclasses))+
  scale_fill_viridis_c(option="magma")


ggsave(plot=last_plot(),filename="Time_to_recovery_lDNA_all_div1.pdf", width = 18, height = 10, units = "cm")


#no effect of N on time to recovery


ggplot(subset(sumstats_median,(as.factor(numGclasses)==4)&(mut==0)), 
       aes(epsilon1,epsilon2,z=tfinal/365))+
  geom_raster(aes(fill = tfinal/365)) +
  #  geom_contour(colour = "white")+
  labs(fill="years to recovery",x=expression(paste(epsilon [1])),y=expression(paste(epsilon [2])))+
  theme(axis.text=element_text(size=10),axis.title=element_text(size=14),plot.title=element_text(size=12,face="bold"))+
  scale_fill_viridis_c(option="magma")


#ggsave(plot=last_plot(),filename="Time_to_recovery_lDNA_N1.pdf", width = 11, height = 8, units = "cm")


#############
#Select runs with the highest fitness (mean)
#############

fraction<-0.025

bestfit1<-sumstats[as.factor(sumstats$numGclasses)==1,] #select runs for a type of class
topTrans1<-quantile(bestfit1$trans,1-fraction) #find the threshold for 10% best fitness
bestfit1<-bestfit1[(bestfit1$trans>=topTrans1),] #select the runs with 10% highest fitness
bestfit1<-as.data.frame(bestfit1)

bestfit2<-sumstats[as.factor(sumstats$numGclasses)==2,] #select runs for a type of class
topTrans2<-quantile(bestfit2$trans,1-fraction) #find the threshold for 10% best fitness
bestfit2<-bestfit2[(bestfit2$trans>=topTrans2),] #select the runs with 10% highest fitness
bestfit2<-as.data.frame(bestfit2)
 
bestfit3<-sumstats[as.factor(sumstats$numGclasses)==3,]#select runs for a type of class
topTrans3<-quantile(bestfit3$trans,1-fraction) #find the threshold for 10% best fitness
bestfit3<-bestfit3[(bestfit3$trans>=topTrans3),] #select the runs with 10% highest fitness
bestfit3<-as.data.frame(bestfit3)


# bestfit5<-sumstats[as.factor(sumstats$numGclasses)==5,] #select runs for a type of class
# topTrans5<-quantile(bestfit5$trans,0.9) #find the threshold for 10% best fitness
# bestfit5<-bestfit5[(bestfit5$trans>=topTrans5),] #select the runs with 10% highest fitness
# bestfit5<-as.data.frame(bestfit5)

bestfit4<-sumstats[as.factor(sumstats$numGclasses)==4,] #select runs for a type of class
topTrans4<-quantile(bestfit4$trans,1-fraction) #find the threshold for 10% best fitness
bestfit4<-bestfit4[(bestfit4$trans>=topTrans4),] #select the runs with 10% highest fitness
bestfit4<-as.data.frame(bestfit4)


bestfit8<-sumstats[as.factor(sumstats$numGclasses)==8,] #select runs for a type of class
topTrans8<-quantile(bestfit8$trans,1-fraction) #find the threshold for 10% best fitness
bestfit8<-bestfit8[(bestfit8$trans>=topTrans8),] #select the runs with 10% highest fitness
bestfit8<-as.data.frame(bestfit8)

bestfit_mean<-rbind(bestfit1,bestfit2,bestfit4,bestfit3,bestfit8)

bestfit_mean$cancer2<-bestfit_mean$cancer>0

bestfit_mean<-as.data.frame(bestfit_mean)


# Mean duration

ggplot(bestfit_mean, aes(x=tfinal/365,colour=as.factor(numGclasses)))+
  geom_density()+ 
  labs(x="Infection duration (in years)",colour="classes")+
  scale_color_viridis_d(option="plasma")+
  theme_light()+
  theme(panel.grid.major = element_blank(),panel.grid.minor = element_blank())+
  geom_vline(data=ddply(bestfit_mean,"numGclasses",summarize,med=mean(tfinal/365)),aes(xintercept=med, colour=as.factor(numGclasses)),linetype="dashed", size=0.5)


ggsave(plot=last_plot(),filename="best_Infection_duration_lDNA.pdf", width = 9, height = 6.5, units = "cm")


ggplot(bestfit_mean, aes(x=tfinal/365,colour=as.factor(cancer>0.0)))+
  geom_density()+ 
  labs(x="Infection duration (in years)",colour="cancer")+
  theme_light()+
  scale_color_grey()+
  facet_wrap(~ as.factor(numGclasses),scale="free")

#ggsave(plot=last_plot(),filename="best_Infection_duration_lDNA_all_div1.pdf", width = 18, height = 10, units = "cm")


# Fitness

ggplot(bestfit_mean, aes(x=log10(trans+1),colour=as.factor(numGclasses)))+
  geom_density()+ 
  labs(x="log10(infection fitness)",colour="classes")+
  theme_light()+
  theme(panel.grid.major = element_blank(),panel.grid.minor = element_blank())+
  scale_color_viridis_d(option="plasma")+
  geom_vline(data=ddply(bestfit_mean,c("numGclasses"),summarize,med=mean(log10(trans+1))),aes(xintercept=med, colour=as.factor(numGclasses)),linetype="dashed", size=0.5)  

  
ggsave(plot=last_plot(),filename="best_fitness_lDNA_all.pdf", width = 9, height = 6.5, units = "cm")


ggplot(bestfit_mean, aes(x=log10(trans+1),colour=as.factor(cancer2)))+
  geom_density()+ 
  labs(x="log10(infection fitness)",colour="cancer > 15%")+
  theme_light()+
  scale_color_grey()+
  facet_wrap(~ as.factor(numGclasses),scale="free")+
  geom_vline(data=ddply(bestfit_mean,c("numGclasses","cancer2"),summarize,med=median(log10(trans+1))),aes(xintercept=med, colour=as.factor(cancer2)),linetype="dashed", size=0.5)  

#ggsave(plot=last_plot(),filename="best_fitness_lDNA_all.pdf", width = 18, height = 10, units = "cm")


# Proportion of cancers

ggplot(bestfit_mean, aes(x=cancer,color=as.factor(numGclasses)))+
  geom_density()+ 
  labs(x="proportion of runs with cancers",colour="classes")+
  theme_light()+
  scale_color_viridis_d(option="plasma")+
  theme(panel.grid.major = element_blank(),panel.grid.minor = element_blank())+
  geom_vline(data=ddply(bestfit_mean,c("numGclasses"),summarize,med=mean(cancer)),aes(xintercept=med, colour=as.factor(numGclasses)),linetype="dashed", size=0.5)  


ggsave(plot=last_plot(),filename="best_proportion_cancers_lDNA_all.pdf", width = 9, height = 6.5, units = "cm")


# Time to cancer

ggplot(subset(bestfit_mean,as.numeric(mut)>0), 
       aes(x=tfinal/365,color=as.factor(numGclasses)))+
  geom_density()+ 
  labs(x="Time to cancer (in years)",colour="classes")+
  scale_color_viridis_d(option="plasma")+
  theme_light() +
  theme(panel.grid.major = element_blank(),panel.grid.minor = element_blank())+
  geom_vline(data=ddply(subset(bestfit_mean,as.numeric(mut)>0),c("numGclasses"),summarize,med=mean(tfinal/365)),aes(xintercept=med, colour=as.factor(numGclasses)),linetype="dashed", size=0.5)  


ggsave(plot=last_plot(),filename="best_Time_to_cancer_lDNA_all.pdf", width = 9, height = 6.5, units = "cm")


####"" epsilon values with or without cancer


ggplot(bestfit_mean, 
       aes(x=epsilon1,colour=as.factor(numGclasses)))+
  geom_density()+ 
  labs(x=expression(paste(epsilon[1])),colour="classes")+
  scale_color_viridis_d(option="plasma")+
  theme_light()+
  theme(panel.grid.major = element_blank(),panel.grid.minor = element_blank())+
  geom_vline(data=ddply(bestfit_mean,c("numGclasses"),summarize,med=mean(epsilon1)),aes(xintercept=med, colour=as.factor(numGclasses)),linetype="dashed", size=0.5)

# 
# 
# ggplot(bestfit_mean, 
#        aes(x=epsilon1,colour=as.factor(cancer2)))+
#   geom_density()+ 
#   labs(x=expression(paste(epsilon[1])),colour="cancer")+
#   scale_color_grey()+
#   theme_light()+
#   facet_wrap(~ as.factor(numGclasses))+
#   geom_vline(data=ddply(bestfit_mean,c("numGclasses","cancer2"),summarize,med=mean(epsilon1)),aes(xintercept=med, colour=as.factor(cancer2)),linetype="dashed", size=0.5)
# 

ggsave(plot=last_plot(),filename="best_epsilon1_lDNA_all.pdf", width = 9, height = 6.5, units = "cm")


ggplot(bestfit_mean, 
       aes(x=epsilon2,colour=as.factor(numGclasses)))+
  geom_density()+ 
  labs(x=expression(paste(epsilon[2])),colour="classes")+
  scale_color_viridis_d(option="plasma")+
  theme_light()+
  theme(panel.grid.major = element_blank(),panel.grid.minor = element_blank())+
  geom_vline(data=ddply(bestfit_mean,c("numGclasses"),summarize,med=mean(epsilon2)),aes(xintercept=med, colour=as.factor(numGclasses)),linetype="dashed", size=0.5)


# ggplot(bestfit_mean,  
#        aes(x=epsilon2,colour=as.factor(cancer>0.0)))+
#   geom_density()+ 
#   labs(x=expression(paste(epsilon[2])),colour="cancer")+
#   scale_color_grey()+
#   theme_light()+
#   facet_wrap(~ as.factor(numGclasses))+
#   geom_vline(data=ddply(bestfit_mean,c("numGclasses","cancer2"),summarize,med=mean(epsilon2)),aes(xintercept=med, colour=as.factor(cancer2)),linetype="dashed", size=0.5)


ggsave(plot=last_plot(),filename="best_epsilon2_lDNA_all.pdf", width = 9, height = 6.5, units = "cm")
```
