## Supplementary material for "Modelling the evolution of viral oncogenesis": R scripts to resimulate the data and generate the plots: plotting_p_effect_retrovir.R

```
library(grid)
library(gridExtra)
library(ggplot2)

#set your working directory
setwd("/home/samuel/Documents/Hote-parasite/HPV/PTRS/Evolution_oncoviruses/Analysis")


###load files
sumstats0<-read.csv(file="sumstats_mean_retrovir0.csv",header = TRUE)
sumstats_median0<-read.csv(file="sumstats_median_retrovir0.csv",header = TRUE)
sumstats0$power<-"p=0"
sumstats_median0$power<-"p=0"
sumstats0$rel_trans<-sumstats0$trans/max(sumstats0$trans)
sumstats_median0$rel_trans<-sumstats_median0$trans/max(sumstats_median0$trans)


sumstats1<-read.csv(file="sumstats_mean_retrovir1.csv",header = TRUE)
sumstats_median1<-read.csv(file="sumstats_median_retrovir1.csv",header = TRUE)
sumstats1$power<-"p=1"
sumstats_median1$power<-"p=1"
sumstats1$rel_trans<-sumstats1$trans/max(sumstats1$trans)
sumstats_median1$rel_trans<-sumstats_median1$trans/max(sumstats_median1$trans)

sumstats2<-read.csv(file="sumstats_mean_retrovir2.csv",header = TRUE)
sumstats_median2<-read.csv(file="sumstats_median_retrovir2.csv",header = TRUE)
sumstats2$power<-"p=2"
sumstats_median2$power<-"p=2"
sumstats2$rel_trans<-sumstats2$trans/max(sumstats2$trans)
sumstats_median2$rel_trans<-sumstats_median2$trans/max(sumstats_median2$trans)

sumstats3<-read.csv(file="sumstats_mean_retrovir3.csv",header = TRUE)
sumstats_median3<-read.csv(file="sumstats_median_retrovir3.csv",header = TRUE)
sumstats3$power<-"p=3"
sumstats_median3$power<-"p=3"
sumstats3$rel_trans<-sumstats3$trans/max(sumstats3$trans)
sumstats_median3$rel_trans<-sumstats_median3$trans/max(sumstats_median3$trans)

sumstats<-rbind(sumstats0,sumstats2,sumstats1,sumstats3)
sumstats_median<-rbind(sumstats_median0,sumstats_median2,sumstats_median1,sumstats_median3)


sumstats<-as.data.frame(sumstats)
sumstats_median<-as.data.frame(sumstats_median)


#############
#Plot Log(infection duration)
#############

ggplot(sumstats, 
       aes(as.numeric(epsilon1),as.numeric(epsilon2),z=as.numeric(log10(tfinal))))+
  geom_raster(aes(fill = as.numeric(log10(tfinal))), interpolate = TRUE) +
  #geom_contour(colour = "white",binwidth = .2)+
  labs(fill=expression(paste(log[10] (T[fin]) )),x=expression(paste(epsilon [1])),y=expression(paste(epsilon [2])))+
  theme(axis.text=element_text(size=10),axis.title=element_text(size=14),plot.title=element_text(size=12,face="bold"))+
  scale_fill_viridis_c(option="inferno")+
  theme_light()+
  theme(legend.title = element_text(size=8))+
  facet_wrap(~ as.factor(power))

ggsave(plot=last_plot(),filename="T_final_retrovir_div.pdf", width = 13, height = 11, units = "cm")


#############
#Plot BH fitness
#############

#for all the number of classes (for comparison)     
ggplot(sumstats, 
       aes(epsilon1,epsilon2,z=log10(trans+1)))+
  geom_raster(aes(fill = log10(trans+1)), interpolate = TRUE) +
# geom_contour(colour = "white")+
  labs(fill="infection\n fitness",x=expression(paste(epsilon [1])),y=expression(paste(epsilon [2])))+
  theme(axis.text=element_text(size=10),axis.title=element_text(size=14),plot.title=element_text(size=12,face="bold"))+
  scale_fill_viridis_c(option="plasma")+
  theme_light()+
  theme(legend.title = element_text(size=8))+
  facet_wrap(~ as.factor(power))


ggsave(plot=last_plot(),filename="BH_Fitness_retrovir_div.pdf", width = 13, height = 11, units = "cm")


ggplot(sumstats, 
       aes(epsilon1,epsilon2,z=100*rel_trans))+
  geom_raster(aes(fill = 100*rel_trans), interpolate = TRUE) +
  # geom_contour(colour = "white")+
  labs(fill="% max trans",x=expression(paste(epsilon [1])),y=expression(paste(epsilon [2])))+
  theme(axis.text=element_text(size=10),axis.title=element_text(size=14),plot.title=element_text(size=12,face="bold"))+
  scale_fill_viridis_c(option="plasma")+
  theme_light()+
  facet_wrap(~ as.factor(power))


#ggsave(plot=last_plot(),filename="relative_BH_Fitness_retrovir_div.pdf", width = 13, height = 11, units = "cm")


#############
#Plot WH fitness
#############

ggplot(sumstats, 
       aes(epsilon1,epsilon2,z=log10(Vtot)))+
  geom_raster(aes(fill = log10(Vtot+1)),interpolate=TRUE) +
  #  geom_contour(colour = "white")+
  labs(fill=expression(paste(log[10] (V[tot]) )),x=expression(paste(epsilon [1])),y=expression(paste(epsilon [2])))+
  theme(axis.text=element_text(size=10),axis.title=element_text(size=14),plot.title=element_text(size=12,face="bold"))+
  scale_fill_viridis_c(option="viridis")+
  theme_light()+
  theme(legend.title = element_text(size=8))+
  facet_wrap(~ as.factor(power))


# N decreases WH fitness (less reinfection)
ggsave(plot=last_plot(),filename="WH_Fitness_retrovir_div.pdf", width = 13, height = 11, units = "cm")


#############
#Plot proportion cancer
#############


ggplot(sumstats, 
       aes(epsilon1,epsilon2,z=cancer))+
  geom_raster(aes(fill = cancer),interpolate=TRUE) +
#  geom_contour(colour = "white")+
  labs(fill=expression(paste(cancer)),x=expression(paste(epsilon [1])),y=expression(paste(epsilon [2])))+
  theme(axis.text=element_text(size=10),axis.title=element_text(size=14),plot.title=element_text(size=12,face="bold"))+
  scale_fill_viridis_c(option="cividis")+
  theme_light()+
  facet_wrap(~ as.factor(power))


ggsave(plot=last_plot(),filename="Cancer_retrovir_div.pdf", width = 13, height = 11, units = "cm")


#############
#Plot time to cancer
#############


ggplot(subset(sumstats,(cancer>0)), 
       aes(epsilon1,epsilon2,z=tfinal/365))+
  geom_raster(aes(fill = tfinal/365),interpolate = TRUE) +
  #  geom_contour(colour = "white")+
  labs(fill="years\n to\n cancer",x=expression(paste(epsilon [1])),y=expression(paste(epsilon [2])))+
  theme(axis.text=element_text(size=10),axis.title=element_text(size=14),plot.title=element_text(size=12,face="bold"))+
  scale_fill_viridis_c(option="magma")+
  theme_light()+
  facet_wrap(~ as.factor(power))


ggsave(plot=last_plot(),filename="Time_to_cancer_retrovir_div.pdf", width = 13, height = 11, units = "cm")


#############
#Plot time to recovery
#############

#for all the number of classes (for comparison)     
ggplot(subset(sumstats,as.numeric(mut)==0), 
       aes(epsilon1,epsilon2,z=tfinal/365))+
  geom_raster(aes(fill = tfinal/365),interpolate = TRUE) +
  #  geom_contour(colour = "white")+
  labs(fill="time to recovery",x=expression(paste(epsilon [1])),y=expression(paste(epsilon [2])))+
  theme(axis.text=element_text(size=10),axis.title=element_text(size=14),plot.title=element_text(size=12,face="bold"))+
  scale_fill_viridis_c(option="magma")+
  theme_light()+
  facet_wrap(~ as.factor(power))


ggsave(plot=last_plot(),filename="Time_to_recovery_retrovir_div.pdf", width = 13, height = 11, units = "cm")


#############
#Select runs with the highest fitness (mean)
#############

fraction<-0.025  #the fraction to keep (here approx 25 runs)
cancer_threshold<-0.0

#topTrans0<-quantile(sumstats0$trans,(1-fraction),na.rm=TRUE) #find the threshold for 10% best fitness
#bestfit0<-sumstats0[(sumstats0$trans>=topTrans0),] #select the runs with 10% highest fitness

topTrans0c<-quantile(sumstats0$trans[sumstats0$cancer>cancer_threshold],(1-fraction),na.rm=TRUE) #fitness for fittest runs with cancer
topTrans0n<-quantile(sumstats0$trans[sumstats0$cancer<=cancer_threshold],(1-fraction),na.rm=TRUE) #fitness for fittest runs without cancer

bestfit0c<-sumstats0[(sumstats0$trans>=topTrans0c)&(sumstats0$cancer>cancer_threshold),] #select the runs with 10% highest fitness
bestfit0n<-sumstats0[(sumstats0$trans>=topTrans0n)&(sumstats0$cancer<=cancer_threshold),] #select the runs with 10% highest fitness

bestfit0<-rbind(bestfit0c,bestfit0n)


#topTrans1<-quantile(sumstats1$trans,(1-fraction),na.rm=TRUE) #find the threshold for 10% best fitness
#bestfit1<-sumstats1[(sumstats1$trans>=topTrans1),] #select the runs with 10% highest fitness

topTrans1c<-quantile(sumstats1$trans[sumstats1$cancer>cancer_threshold],(1-fraction),na.rm=TRUE) #fitness for fittest runs with cancer
topTrans1n<-quantile(sumstats1$trans[sumstats1$cancer<=cancer_threshold],(1-fraction),na.rm=TRUE) #fitness for fittest runs without cancer

bestfit1c<-sumstats1[(sumstats1$trans>=topTrans1c)&(sumstats1$cancer>cancer_threshold),] #select the runs with 10% highest fitness
bestfit1n<-sumstats1[(sumstats1$trans>=topTrans1n)&(sumstats1$cancer<=cancer_threshold),] #select the runs with 10% highest fitness

bestfit1<-rbind(bestfit1c,bestfit1n)


#topTrans2<-quantile(sumstats2$trans,(1-fraction),na.rm=TRUE) #find the threshold for 10% best fitness
#bestfit2<-sumstats2[(sumstats2$trans>=topTrans2),] #select the runs with 10% highest fitness

topTrans2c<-quantile(sumstats2$trans[sumstats2$cancer>cancer_threshold],(1-fraction),na.rm=TRUE) #fitness for fittest runs with cancer
topTrans2n<-quantile(sumstats2$trans[sumstats2$cancer<=cancer_threshold],(1-fraction),na.rm=TRUE) #fitness for fittest runs without cancer

bestfit2c<-sumstats2[(sumstats2$trans>=topTrans2c)&(sumstats2$cancer>cancer_threshold),] #select the runs with 10% highest fitness
bestfit2n<-sumstats2[(sumstats2$trans>=topTrans2n)&(sumstats2$cancer<=cancer_threshold),] #select the runs with 10% highest fitness

bestfit2<-rbind(bestfit2c,bestfit2n)

#topTrans3<-quantile(sumstats3$trans,(1-fraction),na.rm=TRUE) #find the threshold for 10% best fitness
#bestfit3<-sumstats3[(sumstats3$trans>=topTrans3),] #select the runs with 10% highest fitness

topTrans3c<-quantile(sumstats3$trans[sumstats3$cancer>cancer_threshold],(1-fraction),na.rm=TRUE) #fitness for fittest runs with cancer
topTrans3n<-quantile(sumstats3$trans[sumstats3$cancer<=cancer_threshold],(1-fraction),na.rm=TRUE) #fitness for fittest runs without cancer

bestfit3c<-sumstats3[(sumstats3$trans>=topTrans3c)&(sumstats3$cancer>cancer_threshold),] #select the runs with 10% highest fitness
bestfit3n<-sumstats3[(sumstats3$trans>=topTrans3n)&(sumstats3$cancer<=cancer_threshold),] #select the runs with 10% highest fitness

bestfit3<-rbind(bestfit3c,bestfit3n)


bestfit<-rbind(bestfit0,bestfit1,bestfit2,bestfit3)

bestfit$cancer2<-bestfit$cancer>cancer_threshold


bestfit_mean<-as.data.frame(bestfit)


# Mean duration

ggplot(bestfit_mean, aes(x=tfinal/365,colour=power))+
  geom_density()+ 
  labs(x="Infection duration (in years)",colour="power")+
  theme_light()+
  scale_color_viridis_d()+
  geom_vline(data=ddply(bestfit_mean,c("power"),summarize,tfin=mean(tfinal/365)),aes(xintercept=tfin, colour=as.factor(power)),linetype="dashed", size=0.5)+
  
#  facet_wrap(~ as.factor(power),scale="free")


ggsave(plot=last_plot(),filename="best_Infection_duration_retrovir_div.pdf", width = 9, height = 6.5, units = "cm")


# Infection fitness

ggplot(bestfit_mean, aes(x=log10(trans+1),colour=as.factor(cancer2)))+
  geom_density()+ 
  labs(x="log10(infection fitness)",colour="cancer")+
  theme_light()+
  scale_color_grey()+
  geom_vline(data=ddply(bestfit_mean,c("power","cancer2"),summarize,tfin=mean(log10(trans+1))),aes(xintercept=tfin, colour=as.factor(cancer2)),linetype="dashed", size=0.5)+
  facet_wrap(~ as.factor(power),scale="free")

ggsave(plot=last_plot(),filename="best_fitness_retrovir.pdf", width = 13, height = 11, units = "cm")


# Proportion of cancers

ggplot(bestfit_mean, aes(x=cancer,color=power))+
  geom_density()+ 
  labs(x="Proportion of cancers")+
  theme_light()+
  scale_color_viridis_d()+
  geom_vline(data=ddply(bestfit_mean,c("power"),summarize,tfin=mean(cancer)),aes(xintercept=tfin, colour=as.factor(power)),linetype="dashed", size=0.5)
  

ggsave(plot=last_plot(),filename="best_proportion_cancers_retrovir_div.pdf", width = 13, height = 6.5, units = "cm")


# Time to cancer

ggplot(subset(bestfit_mean,cancer2>0), 
       aes(x=tfinal/365,color=power))+
  geom_density()+ 
  labs(x="Time to cancer (in years)")+
  scale_color_viridis_d()+
  theme_light()+
  geom_vline(data=ddply(subset(bestfit_mean,cancer2>0),c("power"),summarize,tfin=mean(tfinal/365)),aes(xintercept=tfin, colour=as.factor(power)),linetype="dashed", size=0.5)


ggsave(plot=last_plot(),filename="best_Time_to_cancer_retrovir_div.pdf", width = 9, height = 6.5, units = "cm")


####"" epsilon values with or without cancer


ggplot(bestfit_mean, 
       aes(x=epsilon1,colour=as.factor(mut>0)))+
  geom_density()+ 
  labs(x=expression(paste(epsilon[1])),colour="cancer")+
  scale_color_grey()+
  theme_light()+
  facet_wrap(~ as.factor(power),scale="free")

ggsave(plot=last_plot(),filename="best_epsilon1_retrovir_div.pdf", width = 13, height = 11, units = "cm")


ggplot(bestfit_mean,  
       aes(x=epsilon2,colour=as.factor(mut>0)))+
  geom_density()+ 
  labs(x=expression(paste(epsilon[2])),colour="cancer")+
  scale_color_grey()+
  theme_light()+
  facet_wrap(~ as.factor(power),scale="free")


ggsave(plot=last_plot(),filename="best_epsilon2_retrovir_div.pdf", width = 13, height = 11, units = "cm")
```
