## Supplementary material for "Modelling the evolution of viral oncogenesis": R scripts to resimulate the data and generate the plots: plotting_waiting_times.R

```
library(grid)
library(gridExtra)
library(ggplot2)

#set your working directory
setwd("/home/samuel/Documents/Hote-parasite/HPV/PTRS/Evolution_oncoviruses/Analysis")


##Generating a distribution of waiting times with different calsses (to clarify things)
dataN1<-rexp(1000,1/1825)
dataN2<-rexp(1000,2/1825)+rexp(1000,2/1825)
dataN3<-rexp(1000,3/1825)+rexp(1000,3/1825)+rexp(1000,3/1825)
dataN4<-rexp(1000,4/1825)+rexp(1000,4/1825)+rexp(1000,4/1825)+rexp(1000,4/1825)
dataN8<-rexp(1000,8/1825)+rexp(1000,8/1825)+rexp(1000,8/1825)+rexp(1000,8/1825)+rexp(1000,8/1825)+rexp(1000,8/1825)+rexp(1000,8/1825)+rexp(1000,8/1825)

data<-c()
data<-rbind(data,cbind(dataN1,rep("1",1000)))
data<-rbind(data,cbind(dataN2,rep("2",1000)))
data<-rbind(data,cbind(dataN3,rep("3",1000)))
data<-rbind(data,cbind(dataN4,rep("4",1000)))
data<-rbind(data,cbind(dataN8,rep("8",1000)))

colnames(data)<-c("years","classes")
data<-as.data.frame(data)
head(data)

data_me<-c()
data_me<-rbind(data_me,c("1",median(dataN1)))
data_me<-rbind(data_me,c("2",median(dataN2)))
data_me<-rbind(data_me,c("3",median(dataN3)))
data_me<-rbind(data_me,c("4",median(dataN4)))
data_me<-rbind(data_me,c("8",median(dataN8)))

colnames(data_me)<-c("classes","years")
data_me<-as.data.frame(data_me)


library(plyr)
ggplot(data,aes(as.numeric(as.character(years))/365,col=as.factor(classes)))+
  geom_density(alpha = 0.1)+
  xlim(0, 10)+
  theme_light()+
  scale_color_viridis_d(option="plasma")+
  labs(x="years",col="G classes") +
  theme(panel.grid.major = element_blank(),panel.grid.minor = element_blank())+
#  geom_vline(data=data_me, aes(xintercept=as.numeric(as.character(data_me$years))/365, colour=as.factor(data_me$classes)),linetype="dashed", size=0.2)
geom_vline(data=ddply(data,c("classes"),summarize,tfin=mean(as.numeric(as.character(years))/365)),aes(xintercept=tfin), size=0.1)+
geom_vline(data=ddply(data,c("classes"),summarize,tfin=median(as.numeric(as.character(years))/365)),aes(xintercept=tfin, colour=as.factor(classes)),linetype="dashed", size=0.5)


ggsave(plot=last_plot(),filename="distribution_waiting_times_lDNA.pdf", width = 9, height = 6.5, units = "cm")
```
