## Supplementary material for "Modelling the evolution of viral oncogenesis": R scripts to resimulate the data and generate the plots: lDNA_2classes_div3.R

```
#setwd("/home/samuel/Documents/Hote-parasite/HPV/PTRS/Evolution_oncoviruses/Gillespie/for_cluster/")


##########################
#load libraries
##########################
library(foreach)
library(doParallel)
#library(tgp)


##########################
#general parameters
##########################
num_cores=6;          #this is the number of cores to use (6 max for cluster)

set.seed(573)         #set seed for random numbers for repeatability


#epsilon 1 values (0-1)
eps1list<- seq(0,1,0.05)  

#epsilon 2 values
eps2list<- seq(0.5,5.1,0.25)

#empty dataframe
psets<-data.frame()

#loop to form 
for (i in 1:length(eps1list)){
  eps1<- eps1list[i]
  for (j in 1:length(eps2list)){
    pset<-data.frame(
      epsilon1 = eps1,
      epsilon2 = eps2list[j],
      nu = 5*10^(-15)) #nu = 10^(-12) - ^(-14)
    psets<-rbind(psets, pset)
  }
}


epscombinations<-dim(psets)[1]  #number of different parameter combinations for (epsilon1,epsilon2)


##########################
#create a cluster with the number of cores
##########################
myCluster <- makeCluster(num_cores, type="PSOCK") 

registerDoParallel(myCluster) #register cluster with foreach package


#create an output file ready to collect the results
write.table( t(c("run","epsilon1","epsilon2","tfinal","G","R","P","Tc","V","mut","trans","Vtot","nu")),  
             file="output_lDNA_2_div3.csv", 
             sep=',', 
             row.names=F, 
             col.names=F )


#jval<-1

##########################
#main loop over all the combinations of epsilon1 and epsilon2
##########################
all_runs_data<-foreach(jval = c(1:epscombinations), .combine = 'rbind') %dopar% 
{

  powerrisk<-3  # if power function: how the number of division increases the risk of carcinogenic mutation
  
  
  numGclasses=as.integer(2);     #number of classes for G to avoid rapid emergence of cancer "by chance"


  runs <- 50           #number of runs per parameter set
  
  
  MaxDiv<-as.integer(30)    #maximum number of cellular divisions
  
  
  
  Tmax <- 18250         #maximum number of days that is 82 years 

    
  ##########################
  #creating an ad hoc function to draw one number in a poisson distribution 
  ##########################
  
  rpoisson<-function(x)
  {
    rpois(1,x)
  }
  
  

    #initiate parameter values (same for all the replicates)
  
  #jval<-1
  
  #from the input of the function
  nu    <-psets[,3][jval]
  epsilon1 <-psets[,1][jval]
  epsilon2 <-psets[,2][jval]
  #
  omega <-0.001   
  delta <-1.0
  sigma <-0.02
  mu    <-0.02
  alpha <- 0.001
  gamma1 <-0.0000001 
  gamma2 <-0.000001 
  K     <-50 
  K2    <-10^(10) #G carrying capacity
  phi   <-2 
  theta <-0.5
  eta   <-numGclasses/1825 #changing of G class towards reactivation every 5/N years on average (div)
  lambda<-1
  
  
  
  ##########################
  # loop over the replicates
  ##########################
  for(irun in c(1:runs))
  {
    
    #print(irun) #print run number
    
    
    #irun<-1
     
    #initialise the variables for this run
    Gvec <- matrix(0,nrow=numGclasses,ncol=MaxDiv)
    Gvec[1,1]<-10
    G<-sum(Gvec)
    Rvec <- matrix(0,nrow=numGclasses,ncol=MaxDiv)
    R  <- sum(Rvec)
    P  <- 0
    Tc <- 1
    V  <- 0
    mut <- rep(0.0,MaxDiv)
    trans<-0
    Vtot<-0
  
    stop<-0
    
    
    #reset the time and time intervals
    t<-0
    
    #initialise results vector
    pop <- c(t,G,R,P,Tc,V,sum(mut),trans,Vtot)
    #popGR<-c(t,Gvec,Rvec,P,Tc,V,C)
    

    
    #time step is per days
    deltat<-0.5
    
    
    #loop until max time reached or extinction
    while((t<=Tmax)&(stop==0))
    {
      
      #how many of each type of events are happenning (drawn from an poisson distribution)
      
      infection      <- if(sum(omega*V*(1-G/K2)*deltat)>0){rpois(1,omega*V*(1-G/K2)*deltat)}else{0};# 
      
      division_result_tot<- if( delta*R*deltat>0){rpois(1, delta*R*deltat)}else{0};
      division_start_tot <- if(sigma*epsilon2*G*(1-G/K2)*deltat>0){rpois(1, sigma*epsilon2*G*(1-G/K2)*deltat)}else{0}; #G has a K
      nat_death_tot   <- if(mu*(1-epsilon1)*G*deltat>0){rpois(1, mu*(1-epsilon1)*G*deltat)}else{0};
      killingG_tot    <- if(alpha*G*Tc*deltat>0){rpois(1, alpha*G*Tc*deltat)}else{0};
      aging_tot       <- if(eta*G*deltat>0){rpois(1, eta*G*deltat)}else{0};
      
      
      #splitting the events accross replication classes
      division_result<-rep(0,numGclasses*MaxDiv)
      if(division_result_tot>0)
      {
        if(division_result_tot<10^4)
        {  
          #find our which categories are non zero
          temp_division_result <- sample(c(1:(MaxDiv*numGclasses)),size=division_result_tot,prob=as.vector(Rvec)/R,replace=T);
          
          #update the long vector
          division_result[as.numeric(names(table(temp_division_result)))]<-table(temp_division_result)
          
          #turn it back into a matrix
          division_result<-matrix(division_result,nrow=numGclasses,ncol=MaxDiv)
        }
        else
        {
          division_result<-round(division_result_tot*Rvec/R)
        }
      }
      else
      {
        division_result<-matrix(division_result,nrow=numGclasses,ncol=MaxDiv)
      }
      
      
      
      division_start<-rep(0,numGclasses*MaxDiv)
      if(division_start_tot>0)
      {
        if(division_start_tot<10^4)
        {
          temp_division_start <- sample(c(1:(MaxDiv*numGclasses)),size=division_start_tot,prob=as.vector(Gvec)/G,replace=T);
          division_start[as.numeric(names(table(temp_division_start)))]<-table(temp_division_start)  
          division_start<-matrix(division_start,nrow=numGclasses,ncol=MaxDiv)
        }
        else
        {
          division_start<-round(division_start_tot*Gvec/G)
        }
      }
      else
      {
        division_start<-matrix(division_start,nrow=numGclasses,ncol=MaxDiv)
      }
      
      nat_death<-rep(0,numGclasses*MaxDiv)
      if(nat_death_tot>0)
      {
        if(nat_death_tot<10^4)
        {
          temp_nat_death <- sample(c(1:(MaxDiv*numGclasses)),size=nat_death_tot,prob=as.vector(Gvec)/G,replace=T);
          nat_death[as.numeric(names(table(temp_nat_death)))]<-table(temp_nat_death)
          nat_death<-matrix(nat_death,nrow=numGclasses,ncol=MaxDiv)
        }
        else
        {
          nat_death<-round(nat_death_tot*Gvec/G)
        }
      }
      else
      {
        nat_death<-matrix(nat_death,nrow=numGclasses,ncol=MaxDiv)
      }
      
      
      killingG<-rep(0,numGclasses*MaxDiv)
      if(killingG_tot>10^4)
      {
        killingG<-round(killingG_tot*Gvec/G)
      }
      else
      {
        if(killingG_tot>0)
        {
          temp_killingG <- sample(c(1:(MaxDiv*numGclasses)),size=killingG_tot,prob=as.vector(Gvec)/G,replace=T);
          killingG[as.numeric(names(table(temp_killingG)))]<-table(temp_killingG)
        }
        
        killingG<-matrix(killingG,nrow=numGclasses,ncol=MaxDiv)
      }
      
      
      
      
      aging<-rep(0,numGclasses*MaxDiv)
      if(aging_tot>10^4)
      {
        aging<-round(aging_tot*Gvec/G)
      }
      else
      {
        if(aging_tot>0)
        {
          temp_aging <- sample(c(1:(MaxDiv*numGclasses)),size=aging_tot,prob=as.vector(Gvec)/G,replace=T);
          aging[as.numeric(names(table(temp_aging)))]<-table(temp_aging)
        }
        
        aging<-matrix(aging,nrow=numGclasses,ncol=MaxDiv)
      }
      
      
      
      #aging       <- if(sum(eta*rowSums(Gvec)*deltat)>0){unlist(lapply(eta*rowSums(Gvec)*deltat,rpoisson))}else{rep(0,numGclasses)};
      
      killingP    <- if(alpha*P*Tc*deltat>0){rpois(1, alpha*P*Tc*deltat)}else{0};
      imm_growthG <- if(gamma1*(G+R)*Tc*(1-Tc/K)*deltat>0){rpois(1, gamma1*(G+R)*Tc*(1-Tc/K)*deltat)}else{0};
      imm_growthP <- if(gamma2*P*Tc*(1-Tc/K)*deltat>0){rpois(1, gamma2*P*Tc*(1-Tc/K)*deltat)}else{0};
      v_production<- if(phi*P*deltat>0){rpois(1, phi*P*deltat)}else{0};
      v_clearance <- if(theta*V*deltat>0){rpois(1, theta*V*deltat)}else{0};
      
  
      mut_process <- if(sum(Gvec)>0){unlist(lapply(nu*colSums(Gvec)*(c(1:MaxDiv)^powerrisk)*deltat,rpoisson))}else{rep(0,MaxDiv)};
      
            
    #  canc_growth <- if(lambda*C*deltat>0){rpois(1, lambda*C*deltat)}else{0};#growth of C
      
      Gvec  <- Gvec + cbind(rep(0,numGclasses),division_result[,-MaxDiv]) - nat_death - division_start - killingG - aging + rbind(rep(0,MaxDiv),aging[-numGclasses,]);
      Gvec[1,1]<-Gvec[1,1]+infection+sum(division_result)  #younger G are produced via infection or via division (assume one of the dividing cells remains old)
      #Gvec[,MaxDiv]  <- Gvec[,MaxDiv] + division_result[,MaxDiv];  #commenting this line out leads to cells not producing offspring once MaxDiv is reached
      Gvec<-Gvec*(Gvec>0) #check that there are no negative numbers
      
      
      Rvec  <- Rvec + division_start - division_result;
      Rvec<-Rvec*(Rvec>0) #check that there are no negative numbers
      
      P  <- max(P + sum(aging[numGclasses,]) - killingP,0);
      Tc <-Tc + imm_growthG + imm_growthP;
      V  <- max(V + v_production - v_clearance,0);
      mut<-mut+ sum(mut_process);

      Vtot<-Vtot + v_production
      
      
      
      #update total number of G cells and R cells    
      G <- sum(Gvec)
      R  <- sum(Rvec)
      
      #update time and store results
      t <- t+deltat
      pop <- rbind(pop,c(t,G,R,P,Tc,V,sum(mut),trans,Vtot))
      
      
      #check whether stopping conditions are met
      stop=(sum(mut)>0)+(G+R<=0);
      
      if(is.na(stop)){stop=100}
      
      
    }
    
    
    
    
    
    ##########################"
    ### Now that the whole time series is run, we add the transmission events
    #########################"
    
    deltaTr<- 60 #every 30 days, there's a contact, transmission possible
    
    #create a matrix with one line per *potential* transmission time
    Ttimes<-c(0:(round(t/deltaTr)-1))*deltaTr
    popTr<-matrix(0,ncol=dim(pop)[[2]],nrow=length(Ttimes))
    popTr[,1]<-Ttimes
    popTr[,8]<-(-1)      #set their Ptransmission to -1 to find them back
    
    #combine the potential transmission matrix to the results matrix
    newpop<-rbind(pop,popTr)
    ordre<-order(newpop[,1])
    newpop2<-newpop[ordre,]
    
    #find our whether there was a transmission for each potential transmission event
    for(ii in c(2:dim(newpop2)[[1]])) #look at the new matrix
    {
      if(newpop2[ii,8]==(-1)) #if we have a line with a potential transmission event
      {
        newpop2[ii,c(2:7,9:dim(pop)[[2]])]<-newpop2[ii-1,c(2:7,9:dim(pop)[[2]])] #update the value at that time by taking the previous time step
        if((newpop2[ii,6])/(10^(4)+newpop2[ii,6])<=0){prob_trans<-0}
        else
        {prob_trans  <- rbinom(1,1,(newpop2[ii,6])/(10^(4)+newpop2[ii,6]));} #determine if there was transmission
        newpop2[ii,8]<-prob_trans; #update the last column accordingly
      }
    }
    
    #we only look at the final line of the output matrix of the run
    output<-newpop2[dim(newpop2)[[1]],]
    
    #we change the transmission event cell so that it's the sum of all the events
    output[8]<-sum(newpop2[,8])
    
    #we store the results for this run 
    write.table( t(c(irun,epsilon1,epsilon2,output,nu)),  
                 file="output_lDNA_2_div3.csv", 
                 append = T, 
                 sep=',', 
                 row.names=F, 
                 col.names=F )
    
  }
  

  
}


stopCluster(myCluster)   #close the cluster
```
