## Supplementary material for "Modelling the evolution of viral oncogenesis": R scripts to resimulate the data and generate the plots: retrovir_div0.R

```
#setwd("/home/samuel/Documents/Hote-parasite/HPV/PTRS/Evolution_oncoviruses/Gillespie")

##########################
#load libraries
##########################
library(tictoc)
library(foreach)
library(doParallel)
#library(tgp)

tic()


##########################
#general parameters
##########################
num_cores=6;          #this is the number of cores to use (6 max for cluster)

set.seed(573)         #set seed for random numbers for repeatability


#epsilon 1 values (0-1)
eps1list<- seq(0,1,0.05)  

#epsilon 2 values
eps2list<- seq(0.5,5.1,0.25)

#empty dataframe
psets<-data.frame()

#loop to form 
for (i in 1:length(eps1list)){
  eps1<- eps1list[i]
  for (j in 1:length(eps2list)){
    pset<-data.frame(
      epsilon1 = eps1,
      epsilon2 = eps2list[j],
      nu = 5*10^(-15)) #nu = 10^(-12) - ^(-14)
    psets<-rbind(psets, pset)
  }
}


epscombinations<-dim(psets)[1]  #number of different parameter combinations for (epsilon1,epsilon2)


# ##########################
# #creating a set of parameters (Latin Hypercube Sampling)
# ##########################
# variables<-rbind(epsilon1=c(0,1),epsilon2=c(0.5,8),nu=c(5*10^(-17),5*10^(-17))) 
# psets<-as.data.frame(lhs(epscombinations,variables,shape=c(1,1,1),mode=c(0,0,0)))
# #plot(psets[,c(1,2)])


##########################
#create a cluster with the number of cores
##########################
myCluster <- makeCluster(num_cores, type="PSOCK") 

registerDoParallel(myCluster) #register cluster with foreach package


#jval<-1

#create an output file ready to collect the results
write.table( t(c("run","epsilon1","epsilon2","tfinal","G","R","Tc","V","mut","trans","Vtot","nu")),  
             file="output_retrovir_div0.csv", 
             sep=',', 
             row.names=F, 
             col.names=F )

##########################
#main loop over all the combinations of epsilon1 and epsilon2
##########################
all_runs_data<-foreach(jval = c(1:epscombinations), .combine = 'rbind') %dopar% 
{
  
  
  powerrisk<-0  # if power function: how the number of division increases the risk of carcinogenic mutation
  

  iruns <- 50           #number of runs per parameter set

  
  MaxDiv<-as.integer(30)    #maximum number of cellular divisions

  
  Tmax <- 18250         #maximum number of days that is 82 years 
  
    
  ##########################
  #creating an ad hoc function to draw one number in a poisson distribution 
  ##########################
  
  rpoisson<-function(x)
  {
    rpois(1,x)
  }
  

  #create an empty vector for the results
  results <- c()#
  
  #
  #initiate parameter values (same for all the replicates)
  omega <-0.001   
  delta <-1.0
  sigma <-0.02
  mu    <-0.02
  alpha <-0.001
  gamma <-0.0000001 
  K     <-50 
  K2    <-10^(10) #G carrying capacity
  phi   <-0.5
  theta <-0.5
  lambda<-1
  
  
  nu    <-psets[,3][jval]#-20
  epsilon1 <-psets[,1][jval]
  epsilon2 <-psets[,2][jval]
  

  ##########################
  # loop over the replicates
  ##########################
  for(irun in c(1:iruns))  
  {
    
    
    #initialise the variables for this run
    
    Gvec <- rep(0.0,MaxDiv)
    Gvec[1]<-10
    G<-sum(Gvec)
    Rvec <- rep(0.0,MaxDiv)
    R  <- sum(Rvec)
    Tc <- 1
    V  <- 0
    P <- 0
    t  <- 0
    mut <- rep(0.0,MaxDiv)
    trans<-0
    Vtot<-0
    
    stop<-0
    
    #initialise results vector
    pop <- c(t,G,R,Tc,V,sum(mut),trans,Vtot)
    
    #time step is 1.0 day
    deltat <- 0.5

        
    #loop until max time reached or extinction
    while((t<=Tmax)&(stop==0))
    {
      
      infection <- if(omega*V*(1-(G+R)/K2)*deltat>0){rpois(1, omega*V*(1-(G+R)/K2)*deltat)}else{0};#G has a K
      
  #    division_result<- if( sum(delta*Rvec*deltat)>0){unlist(lapply(delta*Rvec*deltat,rpoisson))}else{rep(0,MaxDiv)};
  #    division_start <- if(sum(sigma*epsilon2*Gvec*(1-G/K2)*deltat)>0){unlist(lapply(sigma*epsilon2*Gvec*(1-G/K2)*deltat,rpoisson))}else{rep(0,MaxDiv)}; #G has a K
  #    nat_death   <- if(sum(mu*(1-epsilon1)*Gvec*deltat)>0){unlist(lapply(mu*(1-epsilon1)*Gvec*deltat,rpoisson))}else{rep(0,MaxDiv)};
  #    killingG    <- if(sum(alpha*Gvec*Tc*deltat>0)){unlist(lapply(alpha*Gvec*Tc*deltat,rpoisson))}else{rep(0,MaxDiv)};
   
      #calculating the total number of events
      division_result_tot <- if(delta*R*deltat>0){rpois(1, delta*R*deltat)}else{0};
      division_start_tot <- if(sigma*epsilon2*G*(1-G/K2)*deltat>0){rpois(1, sigma*epsilon2*G*(1-G/K2)*deltat)}else{0};#G has a K
      nat_death_tot <- if(mu*(1-epsilon1)*G*deltat>0){rpois(1, mu*(1-epsilon1)*G*deltat)}else{0};
      killingG_tot  <- if(alpha*G*Tc*deltat>0){rpois(1, alpha*G*Tc*deltat)}else{0};

      
      #splitting the events accross replication classes
      division_result<-rep(0,MaxDiv)
      if(division_result_tot>0)
      {
        if(division_result_tot<10^4)
        {
          temp_division_result <- sample(c(1:MaxDiv),size=division_result_tot,prob=Rvec/R,replace=T);
          division_result[as.numeric(names(table(temp_division_result)))]<-table(temp_division_result)  
        }
        else
        {
          # if there are too many cells we can't assign them individually to replication classes and just take proportions
          division_result<-round(division_result_tot*Rvec/R)
        }
      }
      
      division_start<-rep(0,MaxDiv)
      if(division_start_tot>0)
      {
        if(division_start_tot<10^4)
        {
          temp_division_start <- sample(c(1:MaxDiv),size=division_start_tot,prob=Gvec/G,replace=T);
          division_start[as.numeric(names(table(temp_division_start)))]<-table(temp_division_start)
        }
        else
        {
          division_start<-round(division_start_tot*Gvec/G)
        }
      }
      
      nat_death<-rep(0,MaxDiv)
      if(nat_death_tot>0)
      {
        if(nat_death_tot<10^4)
        {
          temp_nat_death <- sample(c(1:MaxDiv),size=nat_death_tot,prob=Gvec/G,replace=T);
          nat_death[as.numeric(names(table(temp_nat_death)))]<-table(temp_nat_death)
        }
        else
        {
          nat_death<-round(nat_death_tot*Gvec/G)
        }
      }
      
      killingG<-rep(0,MaxDiv)
      if(killingG_tot>0)
      {
        if(killingG_tot<10^4)
        {
          temp_killingG <- sample(c(1:MaxDiv),size=killingG_tot,prob=Gvec/G,replace=T);
          killingG[as.numeric(names(table(temp_killingG)))]<-table(temp_killingG)
        }
        else
        {
          killingG<-round(killingG_tot*Gvec/G)
        }
        
      }      
      
      v_production <-if(phi*G*deltat>0){rpois(1, phi*G*deltat)}else{0};
      
      imm_growth<- if(gamma*(G+R)*Tc*(1-Tc/K)*deltat>0){rpois(1, gamma*(G+R)*Tc*(1-Tc/K)*deltat)}else{0};
      v_clearance <- if(theta*V*deltat>0){rpois(1, theta*V*deltat)}else{0};
      
      #mutation process depends on cell age
      #    mut_process <- if(sum(nu*Gvec*(c(1:MaxDiv)^powerrisk)*deltat)>0){unlist(lapply(nu*Gvec*(c(1:MaxDiv)^powerrisk)*deltat,rpoisson))}else{rep(0,MaxDiv)};
      
      mut_process <- if(sum(Gvec)>0){unlist(lapply(nu*Gvec*(c(1:MaxDiv)^powerrisk)*deltat,rpoisson))}else{rep(0,MaxDiv)};
      
      Gvec[1]<-Gvec[1] + infection + sum(division_result);      # new cells born after division are age 0
      Gvec<-Gvec+c(0,division_result[-MaxDiv])                  # mother cells move to the next class 
      #Gvec[MaxDiv]<-Gvec[MaxDiv]+ division_result[MaxDiv]               # if we remove this line, cells with MaxDiv do not produce offspring anymore
      Gvec  <- Gvec - nat_death - division_start - killingG;     # death terms
      Rvec  <- Rvec + division_start - division_result;
      Tc <-Tc + imm_growth;
      V  <- V + v_production - v_clearance;
      mut<-mut+ mut_process;
      Vtot<-Vtot + sum(v_production);
      
      
      #check that numbers are positive for Rvec and Gvec (poisson can do weird stuff)
      Rvec<-Rvec*(Rvec>=0)
      Gvec<-Gvec*(Gvec>=0)
      
      #update total number of G cells and R cells    
      G<-sum(Gvec)
      R  <- sum(Rvec)
      
      #update time and store results
      t <- t+deltat
      pop <- rbind(pop,c(t,G,R,Tc,V,sum(mut),trans,Vtot))
      
      #check whether stopping conditions are met
      stop=(sum(mut)>0)+(G+R<=0);
      
      if(is.na(stop)){stop=100}
      
      #    print(Gvec)    
      #    print(Rvec)
      #    print(stop)
      
    }
    
    
    
    
    
    ##########################"
    ### Now that the whole time series is run, we add the transmission events
    #########################"
    
    
    deltaTr<- 60 #every 30 days, there's a contact, transmission possible
    
    #create a matrix with one line per *potential* transmission time
    Ttimes<-c(0:(round(t/deltaTr)-1))*deltaTr
    popTr<-matrix(0,ncol=dim(pop)[[2]],nrow=length(Ttimes))
    popTr[,1]<-Ttimes
    popTr[,7]<-(-1)      #set their Ptransmission to -1 to find them back
    
    #combine the potential transmission matrix to the results matrix
    newpop<-rbind(pop,popTr)
    ordre<-order(newpop[,1])
    newpop2<-newpop[ordre,]
    
    #find our whether there was a transmission for each potential transmission event
    for(ii in c(1:dim(newpop2)[[1]])) #look at the new matrix
    {
      if(newpop2[ii,7]==(-1)) #if we have a line with a potential transmission event
      {
        newpop2[ii,c(2:6,8:dim(pop)[[2]])]<-newpop2[ii-1,c(2:6,8:dim(pop)[[2]])] #update the value at that time by taking the previous time step
        if((newpop2[ii,5])/(10^(4)+newpop2[ii,5])<=0){prob_trans<-0}
        else
        {prob_trans  <- rbinom(1,1,(newpop2[ii,5])/(10^(4)+newpop2[ii,5]));} #determine if there was transmission
        newpop2[ii,7]<-prob_trans; #update the last column accordingly
      }
    }
    
    #we only look at the final line of the output matrix of the run
    output<-newpop2[dim(newpop2)[[1]],]
    
    #we change the transmission event cell so that it's the sum of all the events
    output[7]<-sum(newpop2[,7])
    
    #we store the results for this run 
    #results <- rbind(results,c(irun,epsilon1,epsilon2,output,nu))
    
    write.table( t(c(irun,epsilon1,epsilon2,output,nu)),  
                 file="output_retrovir_div0.csv", 
                 append = T, 
                 sep=',', 
                 row.names=F, 
                 col.names=F )
    
  }
  
  

  
}


stopCluster(myCluster)   #close the cluster


#save the results to a file
#write.csv(all_runs_data, file = "allruns_retrovir_div.csv",row.names = F)

toc()  #to measure how fast the script runs
```
