## Supplementary material for "Modelling the evolution of viral oncogenesis": R scripts to resimulate the data and generate the plots: retrovir_new_single_run.R

```
setwd("/home/samuel/Documents/Hote-parasite/HPV/PTRS/Evolution_oncoviruses/Gillespie/single_runs")


##########################
#load libraries
##########################
library(tictoc)


tic()

set.seed(578) 

##########################
#creating an ad hoc function to draw one number in a poisson distribution 
##########################


MaxDiv<-as.integer(30)    #maximum number of cellular divisions
powerrisk<-1  # if power function: how the number of division increases the risk of carcinogenic mutation


epsilon1 <- 0.0

epsilon2 <- 1.0


Tmax <- 18250         #maximum number of days that is 50 years 

nu    <- 5*10^(-15) #nu = 10^(-12) - ^(-14)

omega <-0.001   
delta <-1.0
sigma <-0.02
mu    <-0.02
alpha <-0.001
gamma <-0.0000001 
K     <-50 
K2    <-10^(10) #G carrying capacity
phi   <-0.5
theta <-0.5
lambda<-1

iruns<-10

results<-list(length=iruns)
  

for(i in c(1:iruns))  
{
  print(i)
  
  rpoisson<-function(x)
  {
    rpois(1,x)
  }
  
  
  
    #initialise the variables for this run
  
  Gvec <- rep(0.0,MaxDiv)
  Gvec[1]<-10
  G<-sum(Gvec)
  Rvec <- rep(0.0,MaxDiv)
  R  <- sum(Rvec)
  Tc <- 1
  V  <- 0
  P <- 0
  t  <- 0
  mut <- rep(0.0,MaxDiv)
  trans<-0
  Vtot<-0
  
  stop<-0
  
  #initialise results vector
  pop <- c(t,G,R,Tc,V,sum(mut),trans,Vtot)
  
  #time step is 1.0 day
  deltat <- 0.5
  
  
  #loop until max time reached or extinction
  while((t<=Tmax)&(stop==0))
  {
    
    infection <- if(omega*V*(1-(G+R)/K2)*deltat>0){rpois(1, omega*V*(1-(G+R)/K2)*deltat)}else{0};#G has a K
    
    #    division_result<- if( sum(delta*Rvec*deltat)>0){unlist(lapply(delta*Rvec*deltat,rpoisson))}else{rep(0,MaxDiv)};
    #    division_start <- if(sum(sigma*epsilon2*Gvec*(1-G/K2)*deltat)>0){unlist(lapply(sigma*epsilon2*Gvec*(1-G/K2)*deltat,rpoisson))}else{rep(0,MaxDiv)}; #G has a K
    #    nat_death   <- if(sum(mu*(1-epsilon1)*Gvec*deltat)>0){unlist(lapply(mu*(1-epsilon1)*Gvec*deltat,rpoisson))}else{rep(0,MaxDiv)};
    #    killingG    <- if(sum(alpha*Gvec*Tc*deltat>0)){unlist(lapply(alpha*Gvec*Tc*deltat,rpoisson))}else{rep(0,MaxDiv)};
    
    #calculating the total number of events
    division_result_tot <- if(delta*R*deltat>0){rpois(1, delta*R*deltat)}else{0};
    division_start_tot <- if(sigma*epsilon2*G*(1-G/K2)*deltat>0){rpois(1, sigma*epsilon2*G*(1-G/K2)*deltat)}else{0};#G has a K
    nat_death_tot <- if(mu*(1-epsilon1)*G*deltat>0){rpois(1, mu*(1-epsilon1)*G*deltat)}else{0};
    killingG_tot  <- if(alpha*G*Tc*deltat>0){rpois(1, alpha*G*Tc*deltat)}else{0};
    
    
    #splitting the events accross replication classes
    division_result<-rep(0,MaxDiv)
    if(division_result_tot>0)
    {
      if(division_result_tot<10^4)
      {
        temp_division_result <- sample(c(1:MaxDiv),size=division_result_tot,prob=Rvec/R,replace=T);
        division_result[as.numeric(names(table(temp_division_result)))]<-table(temp_division_result)  
      }
      else
      {
        # if there are too many cells we can't assign them individually to replication classes and just take proportions
        division_result<-round(division_result_tot*Rvec/R)
      }
    }
    
    division_start<-rep(0,MaxDiv)
    if(division_start_tot>0)
    {
      if(division_start_tot<10^4)
      {
        temp_division_start <- sample(c(1:MaxDiv),size=division_start_tot,prob=Gvec/G,replace=T);
        division_start[as.numeric(names(table(temp_division_start)))]<-table(temp_division_start)
      }
      else
      {
        division_start<-round(division_start_tot*Gvec/G)
      }
    }
    
    nat_death<-rep(0,MaxDiv)
    if(nat_death_tot>0)
    {
      if(nat_death_tot<10^4)
      {
        temp_nat_death <- sample(c(1:MaxDiv),size=nat_death_tot,prob=Gvec/G,replace=T);
        nat_death[as.numeric(names(table(temp_nat_death)))]<-table(temp_nat_death)
      }
      else
      {
        nat_death<-round(nat_death_tot*Gvec/G)
      }
    }
    
    killingG<-rep(0,MaxDiv)
    if(killingG_tot>0)
    {
      if(killingG_tot<10^4)
      {
        temp_killingG <- sample(c(1:MaxDiv),size=killingG_tot,prob=Gvec/G,replace=T);
        killingG[as.numeric(names(table(temp_killingG)))]<-table(temp_killingG)
      }
      else
      {
        killingG<-round(killingG_tot*Gvec/G)
      }
      
    }      
    
    v_production <-if(phi*G*deltat>0){rpois(1, phi*G*deltat)}else{0};
    
    imm_growth<- if(gamma*(G+R)*Tc*(1-Tc/K)*deltat>0){rpois(1, gamma*(G+R)*Tc*(1-Tc/K)*deltat)}else{0};
    v_clearance <- if(theta*V*deltat>0){rpois(1, theta*V*deltat)}else{0};
    
    #mutation process depends on cell age
    #    mut_process <- if(sum(nu*Gvec*(c(1:MaxDiv)^powerrisk)*deltat)>0){unlist(lapply(nu*Gvec*(c(1:MaxDiv)^powerrisk)*deltat,rpoisson))}else{rep(0,MaxDiv)};
    
    mut_process <- if(sum(Gvec)>0){unlist(lapply(nu*Gvec*(c(1:MaxDiv)^powerrisk)*deltat,rpoisson))}else{rep(0,MaxDiv)};
    
    Gvec[1]<-Gvec[1] + infection + sum(division_result);      # new cells born after division are age 0
    Gvec<-Gvec+c(0,division_result[-MaxDiv])                  # mother cells move to the next class 
    #Gvec[MaxDiv]<-Gvec[MaxDiv]+ division_result[MaxDiv]               # if we remove this line, cells with MaxDiv do not produce offspring anymore
    Gvec  <- Gvec - nat_death - division_start - killingG;     # death terms
    Rvec  <- Rvec + division_start - division_result;
    Tc <-Tc + imm_growth;
    V  <- V + v_production - v_clearance;
    mut<-mut+ mut_process;
    Vtot<-Vtot + sum(v_production);
    
    
    #check that numbers are positive for Rvec and Gvec (poisson can do weird stuff)
    Rvec<-Rvec*(Rvec>=0)
    Gvec<-Gvec*(Gvec>=0)
    
    #update total number of G cells and R cells    
    G<-sum(Gvec)
    R  <- sum(Rvec)
    
    #update time and store results
    t <- t+deltat
    pop <- rbind(pop,c(t,G,R,Tc,V,sum(mut),trans,Vtot))
    
    #check whether stopping conditions are met
    stop=(sum(mut)>0)+(G+R<=0);
    
    if(is.na(stop)){stop=100}
    
#   print(Gvec)    
#    print(Rvec)
#    print(stop)
    
  }

    
colnames(pop) <- c("time","G","R","Tc","V","mut","trans","Vtot")

rownames(pop) <-c()    

results[[i]]<-pop

#print(c(t,mut))

}


plot(rep(-10,length(c(0:round(Tmax/365))))~c(0:round(Tmax/365)),type="l",xlab="time (years)",ylab="log[10](population size +1)",xlim=c(0,30),ylim=c(0,10))
for(j in c(1:iruns))
{
  data<-as.data.frame(results[[j]])
  timeY<-as.numeric((data$time/365))
  lines(log10(1+data$G)~timeY,col=1,lwd=0.2)
  lines(log10(1+data$R)~timeY,col=2,lwd=0.2)
  lines(log10(1+data$V)~timeY,col=3,lwd=0.2)
  lines(log10(1+data$Tc)~timeY,col=4,lwd=0.2)
  abline(v=max(timeY),lty=2)
}

which(data$R<0)

numGclasses<-1

gdata<-c()
vdata<-c()
for(ig in c(1:iruns))
{
  data<-as.data.frame(results[[ig]])
  timeY<-as.numeric(as.character(data$time))
  Gs<-cbind(timeY,log10(1+data$G),"G",numGclasses,ig)
  Rs<-cbind(timeY,log10(1+data$R),"R",numGclasses,ig)
  Vs<-cbind(timeY,log10(1+data$V),"V",numGclasses,ig)
  Tc<-cbind(timeY,log10(1+data$Tc),"Tc",numGclasses,ig)
#  Cs<-cbind(max(timeY),0,"end",numGclasses,ig)
  gdata<-rbind(gdata,Gs,Rs,Vs,Tc)
  if(max(timeY)<Tmax)
  {
  vdata<-rbind(vdata,c(max(timeY),numGclasses,ig))
  }
}

colnames(gdata)<-c("time","popsize","type","N","run")
gdata<-as.data.frame(gdata)


write.csv(gdata, file = "single_run_retrovir_cleared_(0-1).csv",row.names = F,quote=F)

toc()


####"" Plotting age distributions as a function of epsilon1


#write.csv2(data,file="age_distribution_cells.csv",row.names = FALSE)

data<-read.csv2(file="age_distribution_cells.csv",header=TRUE)

#  data<-c()
#  data<-rbind(data,cbind(Gvec01,c(1:30),rep("0.1",30)))
#  data<-rbind(data,cbind(Gvec0,c(1:30),rep("0",30)))
#  data<-rbind(data,cbind(Gvec05,c(1:30),rep("0.5",30)))
#  data<-rbind(data,cbind(Gvec1,c(1:30),rep("1",30)))
# # 
# # names(data)<-c(expression(paste(epsilon [1])),"cells")
# head(data)
# colnames(data)<-c("cells","divisions","epsilon1")
data<-as.data.frame(data)

ggplot(data,aes(y=log10(as.numeric(as.character(cells))),x=as.numeric(as.character(divisions)),col=as.factor(epsilon1)))+
  #  geom_smooth()+
  geom_point()+  
  theme_light()+
  scale_color_viridis_d(option="viridis")+
  labs(y=expression(paste(log [10] (cells))),x="divisions",col=expression(paste(epsilon [1])))

ggsave(plot=last_plot(),filename="age_distribution_retrovir_epsilon1.pdf", width = 9, height = 6.5, units = "cm")
```
